# Systemic effects of melanoma-secreted MIDKINE in the inhibition of dendritic cell differentiation and function

**DOI:** 10.1101/2022.12.28.521901

**Authors:** Xavier Catena, Marta Contreras-Alcalde, Daniela Cerezo-Wallis, Naiara Juan-Larrea, David Olmeda, Guadalupe G. Calvo, Cynthia Mucientes, Sergio Oterino, María S. Soengas

## Abstract

Cutaneous melanomas are a prime example of potentially immunogenic tumors, and as such, ideal targets for immune therapy. These lesions have the largest mutational burden described to date, and accumulate a broad spectrum of post-transcriptional and translational alterations that could conceptually result in a plethora of neoantigens for immune recognition. However, a significant fraction of metastatic melanoma patients is or becomes resistant to current immunotherapeutic agents. How lesions that should represent an inherently hot milieu for immune attack shift into immunologically cold or irresponsive neoplasms is not well understood. Combining cellular systems, mouse models and clinical datasets, here we identify the growth factor Midkine (MDK) as a multipronged blocker of antigen presentation. Mechanistically, we found MDK to repress all main aspects of the maturation, activation and function of dendritic cells, particularly of conventional type 1 (cDC1). These roles of MDK were found to involve primary tumors and lymph nodes, and were traced back to suppressive effects on myeloid precursor cells in the bone marrow. Moreover, MDK shifted the transcriptional profile of DCs towards a tolerogenic state that prevented and bypassed CD8^+^ T cell activation. Blocking MDK enhanced the response to DC-based vaccination and improved the response to immune checkpoint blockade.Together, these data provide insight into how melanomas overcome immune surveillance and support MDK as a target for therapeutic intervention.

## INTRODUCTION

Cutaneous melanomas have puzzled the oncology field for their potential immunogenic features and yet, their inherent ability to bypass immune surveillance^1,2^. Thus, these tumors present the highest mutational burden described to date^3^ and accumulate a variety of post-transcriptional and translational alterations that together, could drive a large set of neoantigens^2,4^. Indeed, immune-based responses are at the heart of spontaneous regressions that have long been reported (albeit occasionally) in melanocytic lesions^5^. Moreover, cutaneous melanomas are among the cancer types with the most effective clinical responses to immune checkpoint blockade^6^. However, over 40% of metastatic melanoma patients are or become resistant to current immunotherapeutic regimens^1,7,8^. How melanomas rewire their immune profiles towards “cold” or irresponsive phenotypes is a matter of intense research^1,9^.

A main hurdle to identify targets for therapeutic intervention in melanoma is the fact that the capacity for metastasis and for immune suppression can be acquired at very early stages of the disease and involve systemic mechanisms^10^. Malignant cells from cutaneous lesions of barely 1 mm in depth can already avoid surveillance mechanisms in the skin, the draining lymph nodes, the bone marrow and distal sites, and ultimately, give rise to metastases, typically in multiple organs^11^. A variety of tumor cell intrinsic and extrinsic mechanisms have been reported to drive this remarkable ability to hide and bypass immune recognition^1^. Puzzlingly, comprehensive proteomic analyses have revealed a complex interplay of pro- and anti-tumorigenic signals coexisting in intricate balances within the melanoma secretome^12^. Recurrent mutations in the oncogene BRAF, loss of the tumor suppressor PTEN or alterations in ß-catenin signaling are some genetic alterations of melanocytic cells that can result in the activation of inflammatory signals, involving for example NF-κB or TGF-ß pathways^2,4^. Microenvironmental cues, can however, shift signaling cascades into tolerogenic phenotypes^13,14^. Conversely, a variety of IFN-associated chemokines and cytokines can favor antigen presentation and mobilize CD8^+^ cytotoxic T to tumor sites, but at the same time, control the expression of immune checkpoint blockers in various immune cells (i.e. CTLA4, PD1, PD1, PDL1 and others)^15^. Determinants of these different immunomodulatory scenarios are not well defined.

Our group has developed “MetAlert” mouse models for whole body imaging of tumor progression in melanoma^9,16^. These animals, together with validation studies in human patient biopsies identified the growth factor MIDKINE (MDK) as potent conditioning factor for premetastatic sites^16^. A key component in the mode of action of MDK was the activation of neolymphangiogenesis, a process with multiple implications in the recruitment and rewiring of the immune system^16^. Subsequently, we demonstrated that macrophages were also targets for the action of MDK^17^. Specifically, we found that MDK rewired the expression profile of macrophages toward tumor-promoting phenotypes^17^. However, inhibition of macrophage function, reduced, but not inhibited, MDK-driven melanoma progression and metastasis. Therefore, we questioned the role of MDK on other immune cell types.

Dissecting the mechanism of action of MDK has basic clinical implications, as this protein may exert pro- or anti-inflammatory roles depending on the context. In addition to melanoma, MDK is expressed in a variety of cancer types such as carcinomas of the lung^18^, breast^19^, liver^20^, prostate^21^, colon^22^ and the gastrointestinal track^23^, as well as in oral squamous cell carcinoma^24^ and glioma^25^. Roles vary in these different pathologies, but rank from the modulation of tumor cell stemness, survival and invasion to the resistance to a variety of chemotherapeutic agents^26^. With respect to immune modulation, MDK has been found to repress natural killer cells (NK)^27^ and as the case for melanoma, promote suppressive macrophage^28^. However, MDK may act in a seemingly opposing manner in other settings, namely, priming, instead of blocking immune responses^29^. In particular, MDK can be induced by pro-inflammatory and anti-tumorigenic cytokines such as IL-2 or IFN-γ^30^. Moreover, MDK has been found linked to a variety of autoimmune and degenerative diseases^29,31,32^. In these settings, MDK can act as a potent recruiter of leukocytes^33^ (i.e., neutrophils, monocytes, macrophages or mast cells, among others) that favor cytotoxic T cell activation^34^. Effects of MDK at lymph nodes and bone marrow that could result in pro vs anti-inflammatory roles are not known.

In this study, we focus on long-range acting roles of MDK on melanoma progression. We dedicated particular emphasis to signaling cascades with the potential for coordinated systemic actions (namely, at lymph nodes and bone marrow, that could also involve immune cells in circulation). Comparative analyses were also performed with respect to other cancer types, to further extent the clinical implication of our findings. These studies were complemented with treatment-based vaccinations and immune checkpoint blockade. This approach revealed an unexpectedly potent inhibitory activity of MDK in antigen presentation, particularly via conventional type I dendritic cells (cDC1). cDC1s are of special interest as they are limited to very low levels in melanoma and other tumor types, but the underlying mechanisms are not well understood. This strategy uncovered new roles of MDK as a signaling hub controlling the maturation, activation and functioning of cDC1, with clinical implications in the resistance to immune-based therapies.

## RESULTS

### Transcriptomic analyses identify MIDKINE as novel regulator of dendritic cell differentiation and function

We have previously reported that MDK depletion promotes large changes in the mRNA and protein expression profiles of cultured melanoma cells^17^. To further define roles of MDK *in vivo*, and possibly, identify new downstream effectors of this protein, we performed a comprehensive characterization of MDK-associated gene signatures using murine models and large human patient datasets. These studies were then combined with functional studies in cellular and mouse models, for validation in the context of vaccination approaches in mice, and of immune checkpoint blockade in various cohorts of melanoma patients. Specifically, we started by a transcriptomic analysis using RNA sequencing (RNAseq) from melanoma allografts generated in immune competent mice (see analyses in human patient datasets below). For tumor implantation, we selected B16.R2L, as an example of a highly metastatic and immunogenic melanoma cell line^35^ that expresses high levels of Mdk^17^ (herein we will use MDK vs Mdk nomenclature to refer to human or murine genes, respectively). These cells were then transduced with lentiviral vectors driving a short hairpin, which we had validated for an efficient and stable depletion of *Mdk* (shMdk). Controls included parental cells expressing non-targeting control shRNA (shCtrl). As summarized in **Fig. 1a**, MDK depletion resulted in broad changes in gene expression *in vivo*. Significantly downregulated genes (n=53) were found with ClueGo (Cytoscape platform)^36^,^37^ to be enriched in developmental growth and epithelial and mesenchymal transition, while upregulated factors (n=362) were primarily enriched in immune modulation and a variety of stress-related programs. A more in-depth analysis of these immune-associated processes upon MDK downregulation revealed programs associated with immune responses, myeloid and leukocyte differentiation (**Fig. 1b**), consistent with our previous data^17^.

**Figure 1.**
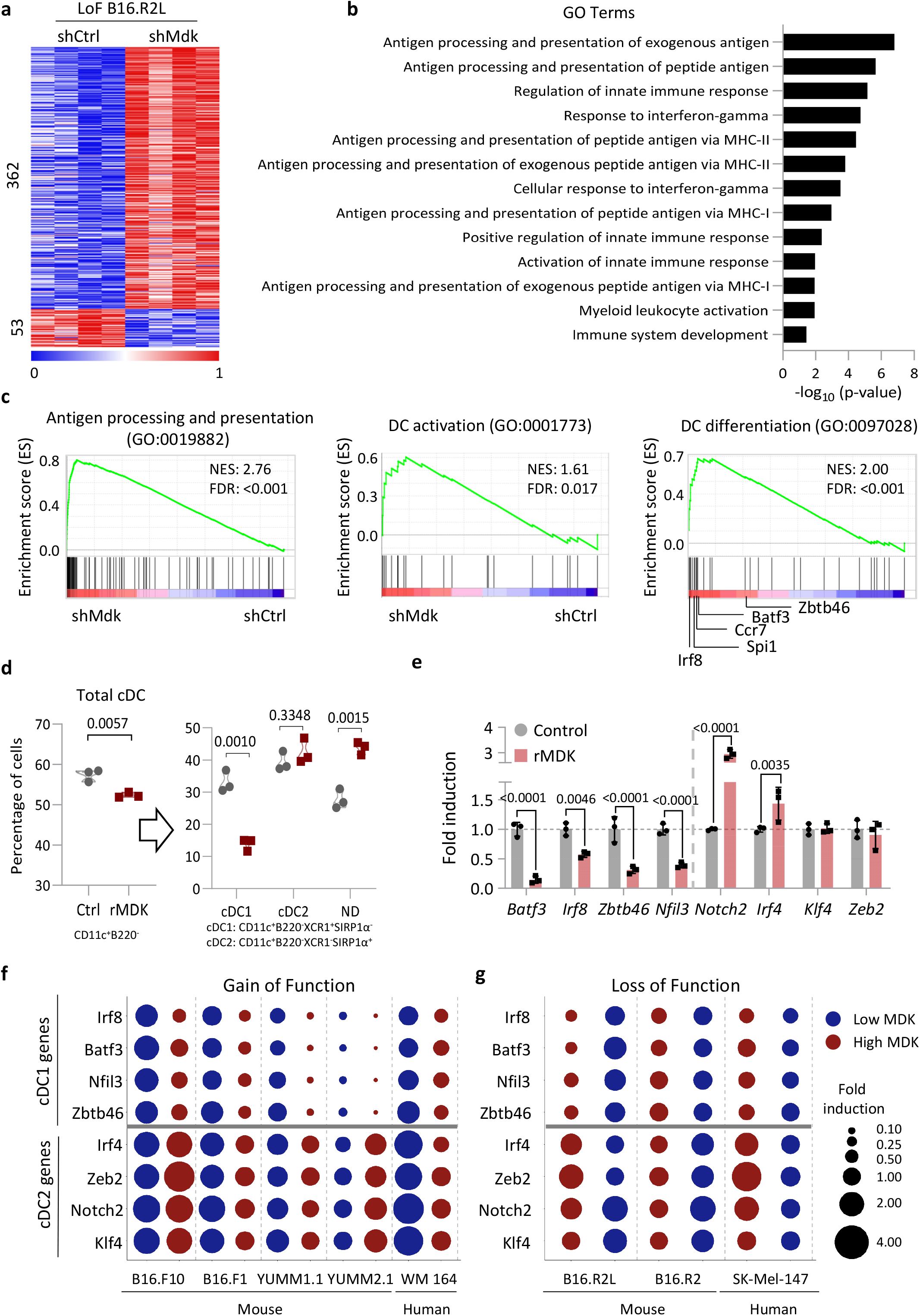
Transcriptomic analyses of melanoma identify new inhibitory roles of MDK in antigen presentation. **(a)** Heatmap of genes differentially expressed in tumor implants generated by B16.R2L cells transduced with lentivirus expressing (shCtrl) or made deficient for MDK (shMdk) and addressed by RNAseq. Scale bar corresponds to reads per kilobase million (RPKM, log_2_ scale) normalized per gene (0 = min, 1 = max). **(b)** Significantly dysregulated immune processes identified using Cytoscape (3.8.2) and ClueGo (2.5.8) from RNAseq data of B16.R2L-shMdk vs shControl allografts generated as in (**a**). Statistical analysis corrected with Bonferroni step down. **(c)** Enrichment scores of the indicated gene ontology (GO) terms defined by GSEA in B16.R2L-shMdk vs shC allografts. **(d)** Percentage of cDCs and cDC1s and cDC2s obtained from FLT3L-incubated BMDCs treated with rMDK (10 ng/ml). Statistical significance was determined by an unpaired two-tailed t-test. **(e)** Relative mRNA levels of the indicated genes defined by qRT-PCR in bone marrow cells cultured with Flt3 (50 ng/ml) and treated with human rMDK (10 ng/ml) during the first three days of the differentiation. Two-way ANOVA corrected for multiple comparisons with Sidak test was used to assess statistical significance. **(f,g)** Relative mRNA expression of the indicated factors in BMDCs incubated with conditional media obtained from the corresponding murine or human cell lines transduced with lentiviral vectors to overexpress (**f**) or deplete MDK (**g**). Circle size represents fold induction with respect to control bone marrow cells (no conditioned media from melanoma cells).

However, and unexpectedly, the most enriched gene ontology (GO) pathways in MDK-depleted allographs were linked to antigen processing and presentation (see **Fig. 1b** and **Fig. 1c**, GO:0019882), most notably signaling cascades associated with dendritic cell (DC) maturation and DC differentiation (**Fig. 1c**, GO:0097028 and GO:0001773). These data suggested a novel double role of MDK controlling differentiation and functional activity of DCs

Genes particularly induced in MDK-depleted allographs were a set of factors such as *Irf8, Spi1, Ccr7, Batf3* and *Zbtb46* (**Fig. 1c, right panel**), characteristically associated with the differentiation of conventional DC1 cells (cDC1)^38^. cDC1s are the most potent antigen presenting subtype of cDCs and their levels are set to particularly low levels in melanoma^13^. We then set to functionally characterize the impact of MDK on this cell type. We also tested for markers and effectors cDC2, another major subset of conventional DCs^38^. To this end, we proceeded to a well-described protocol whereby bone marrow precursors isolated from wild type mice are subjected to incubation with the DC inducer FLT3L^39^. The effect of recombinant MDK (rMDK) was defined by flow cytometry, monitoring total cDCs (CD11c^+^B220^-^), and the effect on cDC1 (CD11c^+^B220^-^XCR1^+^Sirp1a^-^) and cDC2 (CD11c^+^B220^-^XCR1^-^Sirp1a^+^) essentially as described in the Methods section. rMDK reduced the amount of cDCs generated in this protocol, particularly compromising cDC1 (**Fig. 1c**). In the same conditions, cDC2 populations were not significantly affected or a trend was found to increase, not reduce, their expression (**Fig. 1d**). RT-PCR confirmed a differential inhibitory role of human rMDK in transcription factors classically associated to cDC1, but not cDC2 (**Fig. 1e**, see legend for the genes tested. With respect to other DC populations, no significant effects of rMDK were found in plasmacytoid DCs (pDCs, CD11c^+^B220^+^), were found in this system (not shown).

Next, we set to test the effect of MDK on cDC1 vs cDC2 differentiation in a larger panel of melanoma cell lines with endogenously low or high MDK, and modulating the levels of this protein by transducing the corresponding MDK cDNA or shRNAs (i.e gain or loss of function, respectively). As low expressors we used the murine B16.F1 and B16.F10 isogenic lines of the B16 series; the murine YUMM 1.1 and YUMM 2.1, which recapitulate BRAF-driven melanomas^40^, and the human WM-164, which we have found with low metastatic potential^16^. As high expressors, we compared the B16.R2L (isolated from lung metastasis) used above, with the B16.R2 (from lymph node metastasis)^35^, and the human SK-Mel-147 (with a Q61R mutation in NRAS)^41^. As shown in **Fig. 1f,g**, cDC1-associated genes (*Irf8^42,43^, Batf3^44^, Id2^45^*, and *Nfil3^46^*) where decreased in the gain of function setting, while reduced in the loss of function, respectively, with minor changes in cDC2 genes. These results support a crucial effect of this protein on the modulation of cDC1 fate.

### Midkine recapitulates the phenotype of cDC1 deficient mice

To further confirm the relevance of MDK in controlling cDC1s, we performed tumor growth on the *Baft3*-deficient mice. These animals may have various immunological defects as *Batf3* may be expressed in some subtypes of cDC2s, CD4^+^ effector cells and Tregs^47^, but the most prominent being a deficiency in cDC1s^38,44^. Wild type (WT) littermates were included as reference controls. As shown in **Fig. 2a**, Mdk depletion increased survival in WT animals (left panel), consistent with reduced immune suppressed signals when MDK is lost^17^ (note that the growth rates in culture of shCtrl and shMdk-B16R2L cells are similar^17^). Instead, this differential survival was lost in the *Baft3^-/-^* mice (**Fig. 2a, right panel**). Consistent with these results, tumor implants depleted for Mdk were highly inneficient at inducing CD8^+^ T cell infiltration (**Fig. 2b**). Together, these data support a key role of MDK in harnessing conventional type 1 dendritic cells and their consequent effect on CD8^+^ T cell.

**Figure 2.**
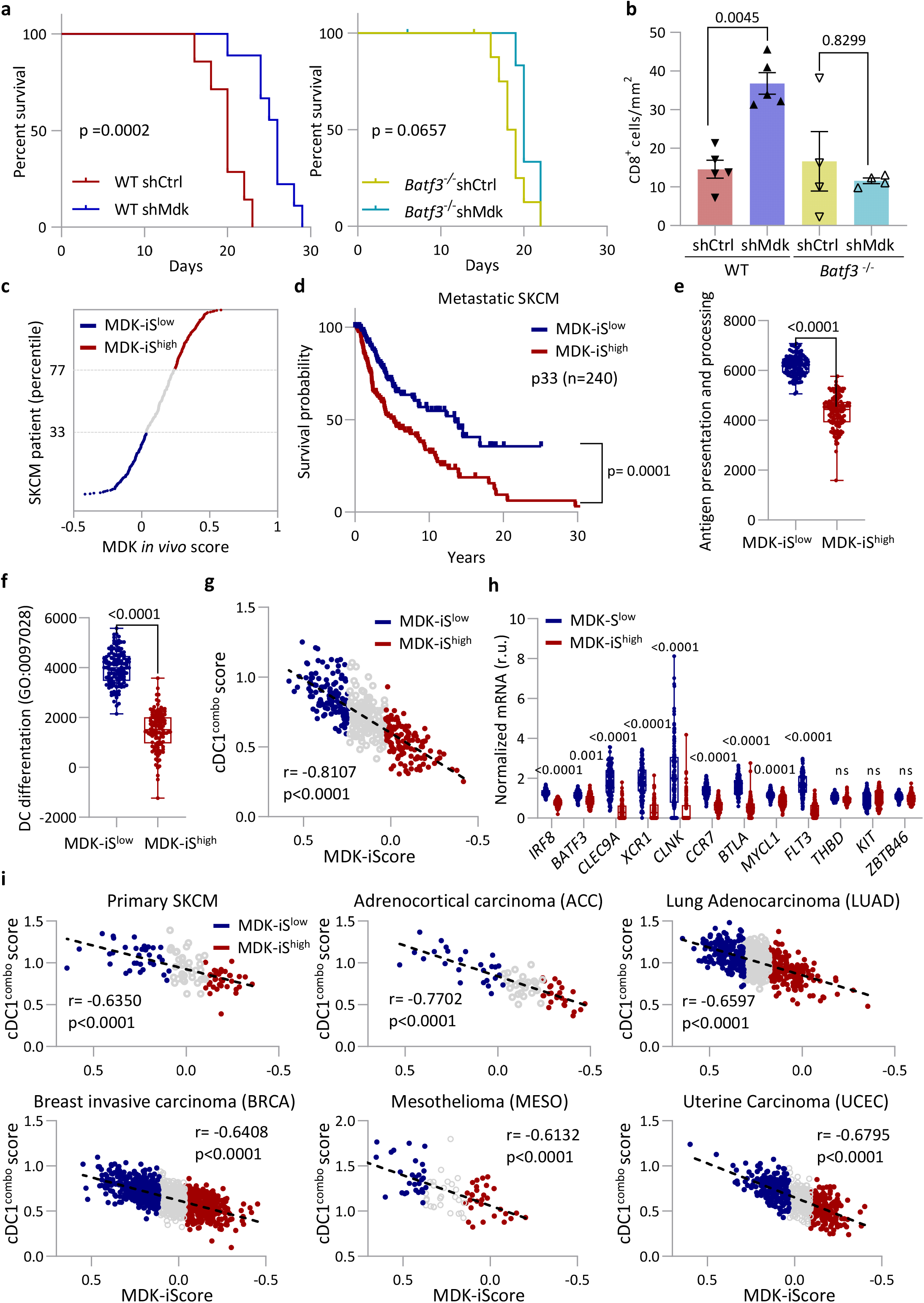
MDK-dependent control of cDC1-associated gene sets. **(a)** Kaplan–Meier curves depicting overall survival of wild type (WT) vs Batf3-deficient mice implanted with B16.R2L-shCtrl or B16R2L-shMdk as indicated. Log-rank (Mantel-Cox) test was performed to compare survival among groups (shown as P value). Samples sizes are n = 7 (WT+shCtrl); n = 9 (WT+shMdk); n = 9 (Batf3^-/-^ +shCtrl); and n = 9 mice (Batf3^-/-^ +shMdk). **(b)** Quantification of intratumoral CD8^+^ T cells by mm^2^ in allografts of B16.R2L (shCtrl vs shMdk) implanted in WT or Batf3^-/-^ mice. One-way ANOVA corrected for multiple comparisons with Tukey test. **(c)** Stratification of TCGA metastatic melanoma patients as a function of the MDK-associated gene score identified in vivo (MDK-iScore) from (Fig. 1a). Patients in the upper and lower third of this scoring are labeled in red and blue, respectively (herein indicated as MDK-iS^high^ and MDK-iS^Low^, respectively). (**d**) Kaplan-Meier curves illustrating the differential overall survival of MDK-iS^high^ vs MDK-iS^Low^ patients in the TCGA melanoma dataset (metastatic lesions). Significance was assessed by Log-rank (Mantel-Cox) test. **(e,f)** Expression of the antigen presentation and processing (GO:0019882), and DC differentiation (GO:0097028) signatures in MDK-iS^low^ and MDK-iS^high^ melanoma patients. Nonparametric unpaired t-test with Mann-Whitney test was applied. **(g)** Correlation of the cDC1^combo^ score and the MDK-iScore in TCGA metastatic melanoma (met-SKCM) patients. Correlation was calculated with the Spearman’s rank correlation coefficient. **(h)** Expression of genes associated to cDC1s in MDK-iS^low^ and MDK-iS^high^ patients. Two-way ANOVA corrected for multiple comparisons with Sidak test was used to assess statistical significance. **(i)** Correlation of cDC1^combo^ score and MDK-iScore within TCGA primary melanoma (primary-SKCM), adrenocortical carcinoma (ACC), lung adenocarcinoma (LUAD), breast invasive carcinoma (BRCA), mesothelioma (MESO) and uterine corpus endometrial carcinoma (UCEC) patients. The Spearman’s rank correlation coefficient was considered to define statistical significance.

### Transcriptomic profiles stratify patients with an inverse MDK-DC expressing score

Next, we turned to patient datasets to define the physiological impact of MDK for patient prognosis and enrichment of DC-associated features. To this end, we took advantage of the well-characterized datasets from The Cancer Genome Atlas (TCGA), which includes metastatic skin cutaneous melanoma (SKCM) patients (n = 364). The top 100 genes we had found upregulated by RNA sequencing on MDK-depleted tumor implants and 44 deregulated ones (**Fig. 1a**) were then used for single sample GSEA in the TCGA SKCM metastatic cohort. This allowed to separate patients as a function of MDK-associated *in vivo* gene score (MDK-iScore, **Fig. 2c**). The top and bottom 1/3 of patients in this analysis, herein referred to as MDK-iS^high^ (n=119) and MDK-iS^low^ (n=121), and depicted in red and blue respectively, in **Fig. 2c**, were selected for further characterization. Importantly, the MDK-iS^high^ patients showed a significantly poorer prognosis than their MDK-iS^low^ counterparts (**Fig. 2d**). Consistent with our data with the depletion of Mdk in mouse allographs, the MDK-iS^low^ showed an enhanced expression of genes involved in antigen presentation and processing (**Fig. 2e**; GO:0019882), and in DC differentiation (**Fig. 2f**; GO:0097028).

As DCs are complex cell populations whose expression profile may vary among systems, we further assessed more specific DC-associated signatures reported in the literature. We thus combined genes that define the so-called Batf3-DC score^48^, a cDC1 signature^49^, and a signature for intratumoral stimulatory dendritic cells (SDC)^50,51^, to create a combined signature we herein refer to as cDC1^combo^ score. Remarkably, this cDC1^combo^ score inversely correlated with the MDK-gene expression profile (**Fig. 2g**, R=0.8107 and p<0.0001). Indeed, mRNA levels of key factors for cDC1 differentiation such as *IRF8, BAFT3, and CLEC9A*, among others, were significantly lower in MDK-iS^high^ than in MDK-iS^low^ patients (**Fig. 2h**).

Of note, the inverse correlation for the MDK-iScore and cDC1^combo^ score was also found to be highly significant in TCGA datasets from primary skin melanoma, adenocortical carcinoma (ADCC), lung adenocarcinoma (LUAD), breast invasive carcinoma (BRCA), mesothelioma (MESO) and uterine corpus endometrial carcinoma (UCEC) (**Fig. 2e**). Together, these data point to new inhibitory effects on DCs in melanoma, which may extend to other tumor types.

### Tumor-driven MDK impacts on DC content at tumor, lymph nodes, blood and bone marrow

Once having defined correlations between MDK levels and cDCs (particularly cDC1), we turned to mouse allografts to address this interplay functionally. First, GoF assays were performed by introducing Mdk in the low expressor B16.F1. Overexpression of MDK in B16.F1 resulted in a reduction of intratumoral cDCs (CD11c^+^MHCII^+^) and cDC1s (CD24^+^ CD103^+^ CD11b^-^), with no significant effects on cDC2 (CD24^-^CD11b^+^CD103^+^), as defined by flow cytometry (**Fig. 3a**; **see Supplementary Fig. 1a-f** for gating strategies used *in vivo*). We then further focused on cDC1s in LoF settings (Fig. **3b-h**; see for cDC2s below in **Supplementary Fig. 1g-j**). Depleting Mdk in the high expressors B16.R2 and B16.R2L resulted in a marked enhancement of intratumoral cDCs and cDC1s (**Fig. 3b-c**). Reduced intratumoral cDCs were also observed by immunofluorescence co-staining for CD11c (green) and MHCII (red) in B16.R2L implants (**Fig. 3d**).

**Figure 3.**
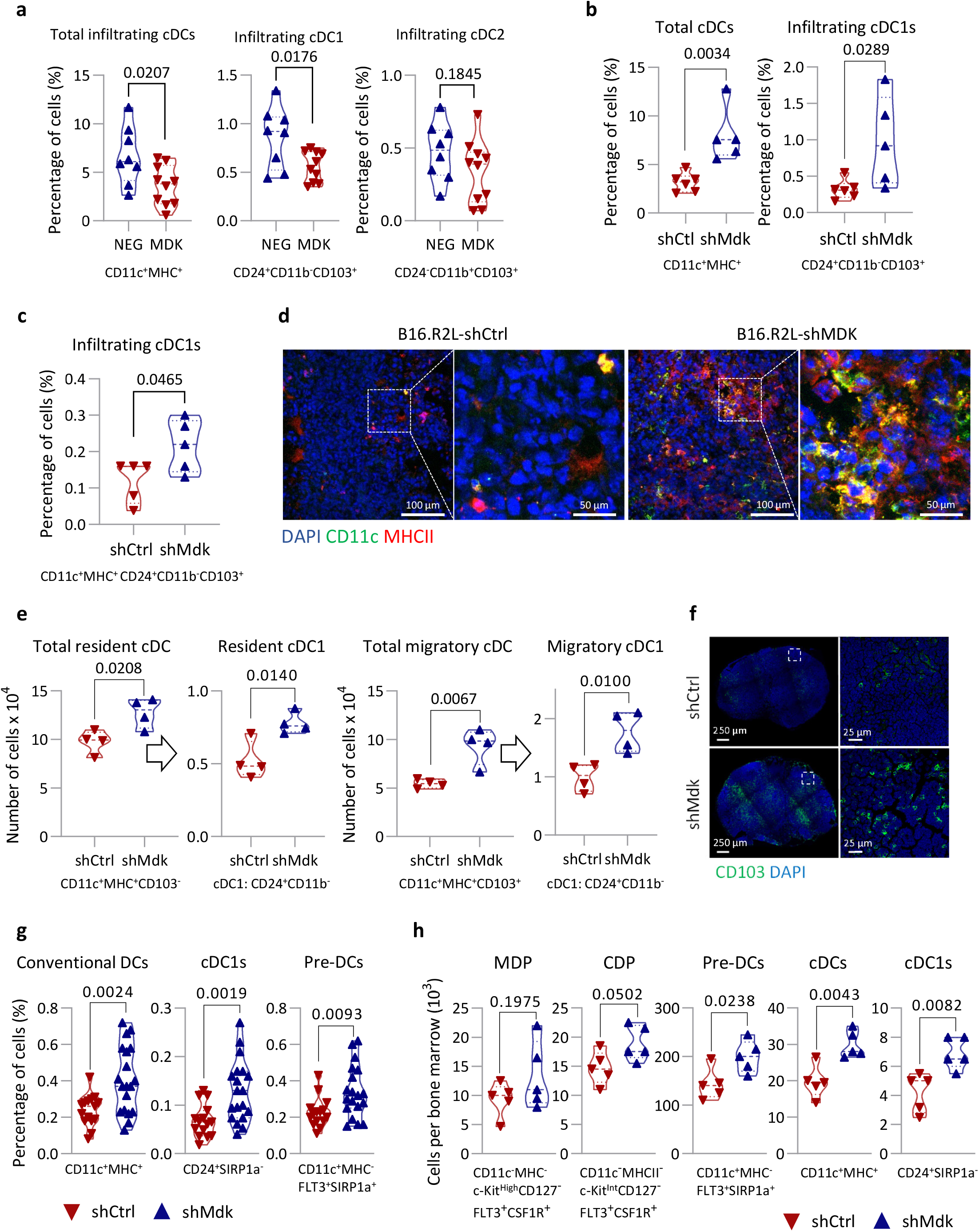
Systemic effects of MDK on dendritic cell populations. **(a)** Infiltration of cDC (CD11c^+^MHCII^+^), cDC1 (CD24^+^CD103^+^), and cDC2 (CD24^-^CD11b^+^CD103^+^) in tumors implanted in immunocompetent mice with B16.F1-MDK cells, as estimated by flow cytometry with respect to their isogenic controls (MDK: n = 10 tumors; Ctrl: n = 8). Statistical significance was determined by an unpaired two-tailed t-test. **(b)** Infiltrating cDC (CD11c^+^MHCII^+^) and cDC1 (CD24^+^CD103^+^) in B16.R2 implants expressing shControl or shMdk (shCtrl: n = 6; shMdk: n =5). Statistical significance was determined by unpaired two-tailed t-test. **(c)** Infiltration of cDC1 (CD24^+^CD103^+^) in B16.R2L-shCtrl and -shMdk melanomas (shCtrl: n = 5; shMdk: n = 5 tumors). Statistical significance was determined by unpaired two-tailed t-test. **(d)** Visualization by immunofluorescence of cDC (CD11c^+^MHCII^+^: yellow) in B16.R2L-shCtrl and shMDK allografts; DAPI in blue, CD11c in green, and MHCII in red. **(e)** Number of DCs in draining lymph nodes (shCtrl: n = 4; shMdk: n = 4 dLN) of B16.R2L-shCtrl and -shMdk melanomas, with the indicated cell types defined by flow cytometry as follows: total resident cDC (CD11c^+^MHCII^+^C103^-^)+cDC1 (CD24^+^CD11b^-^); total migratory cDCs (CD11c^+^MHCII^+^CD103^+^) and cDC1 (CD24^+^CD11b^-^). Statistical significance was determined by unpaired two-tailed t-test. **(f)** Draining lymph nodes from B16.R2L-shCtrl and -shMdk stained with CD103 (migratory DCs in green) and DAPI (blue). **(g)** Percentage of pre-DCs, cDCs, and cDC1s in blood in mice bearing B16.R2L-shCtrl (red) or shMdk (blue) (shCtrl: n = 16; shMdk: n =19). Statistical significance was determined by unpaired two-tailed t-test. **(h)** Number of progenitor and mature dendritic cells in bone marrow preparations (femur and tibia) from mice bearing B16.R2L transduced with shCtrl (red) or shMDK (blue) (shCtrl: n = 5; shMdk: n =5). Macrophage/dendritic cell progenitors (MDPs), common dendritic cell progenitor (CDP), pre-DCs and mature DCs and subtype cDC1 were defined as indicated in the text (see Supplementary Fig. 1). Statistical significance was determined by unpaired two-tailed t-test.

Mdk is a secreted factor that can reach the lymph nodes and distal organs, as we have previously shown. Therefore, we set to assess the effect of Mdk expressed from tumor cells on DC populations beyond the tumor sites. In the lymph nodes, we tested resident cDCs (CD103^-^ and CCR7^-^) and migratory DCs (typically defined by the markers CD103 or CCR7)^52^. As shown in **Fig. 3e-f**, draining lymph nodes of mice implanted with B16.R2L depleted for Mdk showed a higher overall cDC count than the controls, with an increase of both, resident cDCs and migratory CD103^+^ DCs, of the cDC1s subtypes (see by flow cytometry in **Fig. 3e**). Increased DC103^+^ cells in draining lymph nodes of shMdk implants was also observed by immunofluorescence (**Fig. 3f**). Consistent with the mobilization of cDCs, Mdk depletion in B16.R2L resulted also in an enhancement of circulating cDCs and cDC1s (**Fig. 3g**). We also found premature DCs (pre-DCs: CD11c^+^MHC^-^FLT3^+^SIRP1a^+^)^52^ enhanced in the absence of Mdk (**Fig. 3g**).

Considering the data above, we hypothesized that MDK could act at an earlier stage of DC differentiation in the bone marrow. To this end, we analyzed common myeloid progenitors (CMPs), macrophage/dendritic cell progenitors (MDPs), common dendritic cell progenitors (CDPs) in bone marrow preparations from mice implanted with B16.R2L cells expressing or lacking Mdk (pre-DCs and mature cDCs were also analyzed in the same setting). Time points were selected to proceed to flow cytometry analyses before tumor cell dissemination to lymph nodes or distal sites (data not shown). Mice implanted with shMdk-B16.R2L presented no significant changes in common myeloid progenitors (CMPs) (data not shown) and macrophage/dendritic cell progenitors (MDPs) (**Fig. 3h**). Instead, the earlier changes in the DC differentiation cascade were found at the level of common dendritic cell progenitors (CDP, ~30% increase) in addition to pre-DCs (~40%) (**Fig. 3h**). In addition, Mice implanted with shMdk-B16.R2L presented an increase in pre-DCs (~40%), mature cDCs (~50%), cDC1s (~60%) and cDC2s (~45%) (**Fig. 3h**).

Effects of MDK LoF in cDC2s at tumor, lymph node, blood and bone marrow were more variable (**Supplementary Fig. 1g-j**). Therefore, while we further analyzed functional roles of MDK on cDC2s, our prime interest was on cDC1s, based also in data in patients (**Fig. 2c-h**) and as defined using Baft3 deficient mice (**Fig. 2a,b**).

### MDK represses the expression of cDC1-associated transcription factors

Next, we proceeded to profile the impact of MDK on the transriptome of bone marrow derived DCs (BMDCs). For this purpose, murine-extracted bone marrow cells were incubated with the DC-differentiation factor FLT3L (50 ng/ml, 3 days) with or without rMDK (10 ng/ml). rMDK led to marked changes in mRNA expression of BMDCs (458 upregulated and 226 downregulated genes, see heatmaps in **Fig. 4a**). Importantly, as shown in **Fig. 4b**, MDK repressed key genes that define the Batf3-DC score^48^, the cDC1 signature^49^, and the SDC signature^51,53^ we had tested in human patients (**Fig. 2g**). However, this was not the case for cDC2-associated genes^54^ (**Fig. 4b**). As these data were performed in tissue culture, we also addressed expression profiles of bone marrow cells collected early after implantation of B16.R2L-shCtrl (high MDK) or B16.R2L-shMdk (low MDK) in mice. This confirmed a significant induction of cDC1-associated factors (**Fig. 4c**). cDC2 and monocyte-associated factors were instead downregulated (**Fig. 4c**).

**Figure 4.**
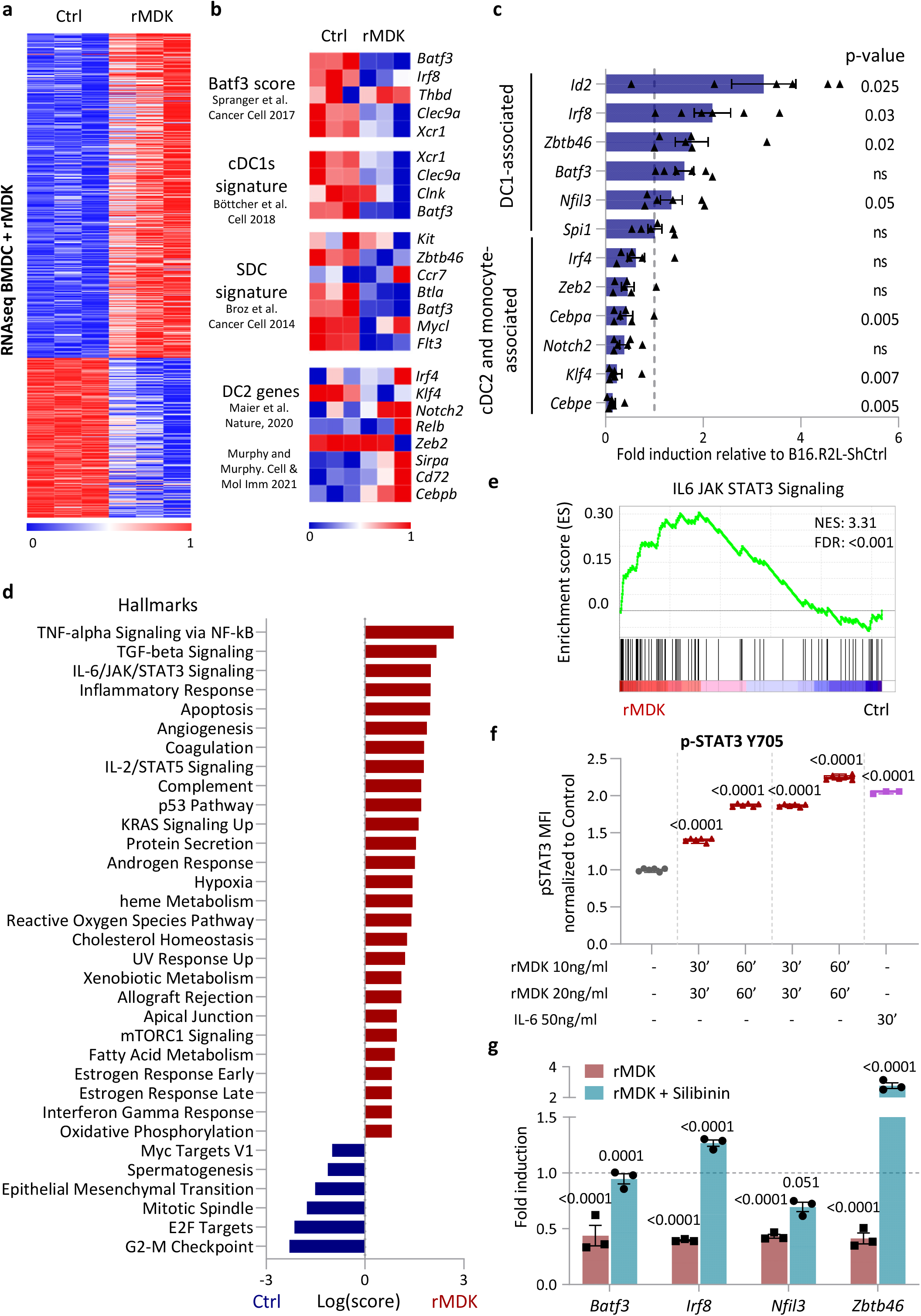
MDK-induced transcriptomic rewiring of DC cells. **(a)** Heatmap of transcriptional changes (RNAseq) obtained upon incubating bone marrow cells with FLT3 (50 ng/ml) in the absence or in the presence of rMDK (10 ng/ml) for 3 days. Scale bar corresponds to reads per kilobase million (RPKM, log_2_ scale) normalized per gene (0 = min, 1 = max). **(b)** Heatmap representing differentially expressed genes in three independent cDC1 signatures and a compendium of cDC2-associated genes^48–51,54^. Data were obtained from transcriptomic analyses of FLT3-bone marrow cultures 3 days after incubation with rMDK. Scale bar corresponds to reads per kilobase million (RPKM, log_2_ scale) normalized per gene (0 = min, 1 = max). **(c)** qRT-PCR analysis of transcription factors associated with dendritic cell and monocyte differentiation expressed by whole bone marrow isolated from mice implanted with B16.R2L-sh*Mdk* (blue), represented with respect to shCtrl controls (dash-line). Statistical significance was obtained by two-way ANOVA corrected for multiple comparisons with Sidak test. **(d)** Enrichment analysis of the indicated pathways (Enrichr-MSigDB Hallmark 2020) in bone marrow cells cultured with FLT3 (50 ng/ml) +/− rMDK (10 ng/ml) for 3 days. **(e)** Gene set enrichment analysis (GSEA) of the IL-6/JAK/STAT3 pathway in FLT3-BMDCs in (d). NES, normalized enrichment score; FDR, False discovery rate. **(f)** Normalized p-STAT3 Y705 defined in DCs at 1, 5, 15 and 30 min after addition of rMDK (10 and 20ng/ml). DCs treated with IL-6 (50 ng/ml) for 30 min was used as a positive control of p-STAT3. One-way ANOVA corrected for multiple comparisons with Tukey test. **(g)** qRT-PCR to assess the expression of Batf3, Irf8, Nfil3 and Zbtb46 in bone marrow cells incubated with the STAT3 inhibitor (Silibinin; 100 μM) 3h before treatment with rMDK (10ng/ml) for 24h. Dash-line represents expression of genes in control cells (no treated with rMDK). Statistical significance was obtained by two-way ANOVA corrected for multiple comparisons with Sidak test.

Next, we set to define more in detail the mRNA expression profiles controlled by MDK in the bone marrow-derived DCs. Enrichment analyses showed that the top upregulated pathway induced by MDK were involved in TNFa, TGFb and IL-6/JAK/STAT3 pathways (**Fig. 4d**). The high enrichment in the IL-6/JAK/STAT3 signaling cascade (**Fig. 4e**; normalized enrichment score of 3.31, FDR < 0.001) was particularly relevant for its known repressive effect of STAT3 on DC^55^, not previously linked to MDK.

Flow cytometry confirmed a dose- and time-dependent phosphorylation of STAT3 in BMDCs by recombinant MDK (see quantification in **Fig. 4f** with respect to IL-6 as a positive reference control). We then tested the impact of the STAT3 inhibitor silibinin on the ability of MDK to repress cDC1-associated transcription factors. As shown in **Fig. 4g,** silibinin significantly compromised the inhibition of *Batf3, Irf8, Nfil3* and *Zbtb46* by MDK. These results reveal a new immune modulatory role of MDK exerted via STAT3-driven repression of key transcription factors involved in DC maturation.

### MDK impairs DC function, which leads to a deficient T cell activation and reduced T cell-mediated melanoma killing

Next, we questioned whether MDK not only compromised the levels of mature DCs, but also their functionality. To this end, we pursued a comprehensive characterization of DCs in phagocytosis, antigen processing, expression of co-stimulatory and co-inhibitory signals and cytokines, lymph node migration, and ultimately, antigen presentation and T cell activation (see schematic of experimental procedures in **Supplementary Fig. 2a**).

A key feature of DCs as antigen-presenting cells (APCs) is their ability to endocyte and phagocyte self and foreign antigens^38^. We then tested the endocytic activity of mature BMDCs by monitoring the uptake of fluorescently labeled ovalbumin (OVA), which was coupled to the AF529 fluorophore. As shown by flow cytometry in **Supplementary Fig. 2b**, rMDK markedly reduced endocytosis of OVA-AF529 particles (see quantification in **Supplementary Fig. 2c**). In addition, we assayed the phagocytosis of mature BMDCs by interrogating the incorporation of *E. coli* BioParticles conjugated to the pHRodo, a pH-sensitive dye that emits green fluorescent in acidic environments, such as in the endosomes or phagosomes. In this case, rMDK was also highly inhibitory (**Fig. 5a**). Indeed, rMDK was as potent as the phagocytic inhibitor cytochalasin D^56^ (see time-lapse analyses depicted in **Fig. 5b**).

**Figure 5.**
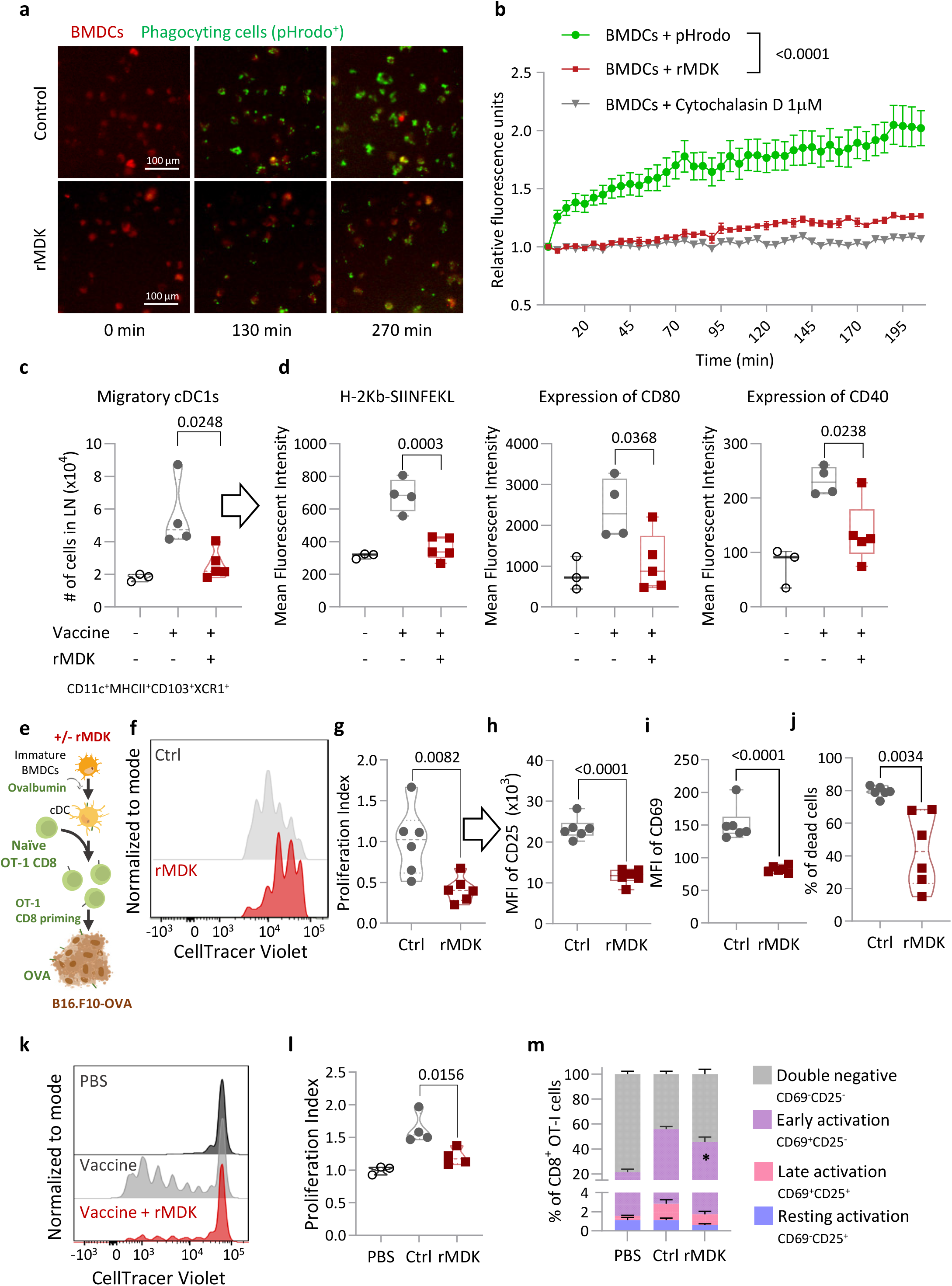
MDK impairs dendritic cell function leading to deficient T cell activation. **(a)** Representative immunofluorescence images of BMDCs phagocyting pHRodo™ E. coli BioParticles™ (green) at three different time points (0 min, 133 min and 273 min), in BMDCs stained with DiD’ (red). **(b)** Quantification of phagocytosis as the relative fluorescence emission per cell estimated with respect to the average levels at the starting point. BMDCs control (green), BMDCs-rMDK (red), BMDCs negative control with Cytochalasin D (grey) upon incubation with pHRodo™ E. coli BioParticles™. Statistical significance was determined by two-way ANOVA corrected for multiple comparisons with Sidak test. **(c)** Number of migratory cDC1s (CD11c^+^MHCII^+^CD24^+^CD11b^-^CD103^+^) in draining lymph nodes from the indicated vaccination arms. Statistical significance assessed by one-way ANOVA corrected for multiple comparisons with Tukey test. **(d)** Expression levels of H-2Kb-SIINFEKL, CD80 and CD40 represented by mean fluorescence intensity (MFI). Statistical significance assessed by one-way ANOVA corrected for multiple comparisons with Tukey test. **(e)** Summary of the experimental approach to define the impact of MDK on DC-mediated antigen priming to CD8+ T cells. **(f)** Representative histogram of *in vitro* OT-I T cell proliferation of BMDCs control (grey) or treated with rMDK for 48h (red) as defined by staining with CellTracer Violet. **(g)** Proliferation index of OT-I T cells primed by BMDCs, either untreated (grey) or treated with rMDK for 48h (red). Data are represented with respect to untreated controls. Statistical significance was determined by unpaired two-tailed t-test. (**h-i**) Expression of CD25 **(h)** and CD69 **(i)** on proliferating OT-I defined by mean fluorescence intensity (MFI). Statistical significance was determined by unpaired two-tailed t-test. **(j)** Percentage of dead B16.F10-OVA melanoma cells in the presence of OT-I T cell primed with BMDCs in the absence (grey) or the presence of rMDK (red). Statistical significance was determined by unpaired two-tailed t-test. **(k)** Representative histogram and **(l)** quantification of OT-I T cell proliferation in vivo or in a vaccination protocol poly(I:C) and CD40) in the absence (grey) or in the presence of rMDK (red). One-way ANOVA corrected for multiple comparisons with Tukey test. **(m)** Evaluation of proliferating OT-I activation by CD69 and CD25 markers (Early activated: CD69^+^CD25^-^; late activated CD69^+^CD25^+^; resting (CD69^-^CD25^+^) and double-negative (DN): CD69^-^CD25^-^). Statistical significance assessed by one-way ANOVA corrected for multiple comparisons with Tukey test, p-value; * =p<0.013.

Next, we assessed the subsequent steps after antigen uptake by DCs, namely, activation to favor antigen presentation. This was then analyzed by monitoring surface expression of MHCI and MHCII molecules in BMDCs, and the induction of co-stimulatory signals (i.e associated with CD40), crucial for T cell activation^38^. Interestingly, activated BMDCs (with LPS) treated with rMDK and pulsed with OVA expressed reduced levels of MHCI, MHCII, and CD40 (**Supplementary Fig. 2d-f**). We also found that rMDK compromised the expression of MHCI loaded with the ovalbumin peptide SIINFEKL (**Supplementary Fig. 2g**).

Once defined a defective activation of DCs *in vitro*, we proceeded to assess this function *in vivo*, using a standard vaccination setting. Briefly, mice were inoculated with a vaccine containing OVA, and the adjuvants poly(I:C) and anti-CD40. Half of these vaccinations included rMDK. 3 days afterwards (time for activated DCs to travel to the draining lymph node and prime T cells), mice were sacrificed and draining lymph nodes were assessed by flow cytometry. As shown in **Fig. 5c**, addition of rMDK to the vaccine reduced the total number migratory cDC1s (CD103^+^XCR1^+^). The number of resident DCs in these lymph nodes remained unchanged (not shown). Consistent with the results in cultured cells (**Supplementary Fig. 2g**), migratory cDC1s from mice vaccinated with rMDK expressed lower levels of the ovalbumin SIINFEKL peptide loaded onto MHCI, as well as a marked reduction of the costimulatory signals CD80 and CD40 (**Fig. 5d**). Together, these results therefore support that MDK reduced DC-dependent antigen presentation and activation in response to vaccination protocols.

A determinant function of cDC1s is their ability to present antigens to CD8^+^ T cells. To asses this process, we used OT-I CD8^+^ T cells. These are transgenic CD8^+^ T cells that specifically recognize the OVA-derived peptide SIINFEKL^57^. Once DCs present this OVA-peptide to OT-I CD8^+^ T cells, they divide and get activated, first expressing the CD69 (early activation) marker and then becoming CD69-CD25 double positive (late activation)^38^. To follow these processes, DCs were incubated with ovalbumin in the presence or absence of MDK. 24h later, these MDK-conditioned DCs were placed in contact with naïve OT-I CD8^+^ T cells labeled with CellTrace Violet. Next day, OT-I CD8^+^ T cell proliferation was monitored by flow cytometry by means of the dilution of the CellTrace Violet signal (**Fig. 5e**). Flow cytometry was also used to assess CD69 and CD25 levels in OT-I cells. Interestingly, OT-I cells showed nearly a 65% reduction of cell proliferation (**Fig. 5f-g**) and reduced by half the expression of CD69 and CD25 (**Fig. 5h-i**) when they were incubated with MDK-conditioned DCs (see p-values in the corresponding figure panels). Finally, we tested the effect of the priming activity of MDK-educated DCs on the cytotoxic activity of OT-I CD8^+^ T. As shown in **Fig. 5j**, OT-I T cells primed by_MDK-DCs presented a reduced 50% melanoma cell killing with respect to the OT-I T cells primed DC controls.

To validate the impact of MDK on antigen presentation *in vivo*, mice were vaccinated with a cocktail of OVA, Poly(I:C) and CD40 agonist in the presence or absence of rMDK. 24 h afterwards, CellTrace Violet-stained OT-I naïve T cells were inoculated retro-orbitally. Three days later, draining lymph nodes were harvested and T cell proliferation, and activation were determined by flow cytometry. In this setting, vaccines containing rMDK showed a reduced ability to promote OT-I CD8^+^ T cell proliferation (**Fig. 5k,l**) and this population presented lower numbers of cells in an early activated phenotype (CD69^+^CD25^-^; **Fig. 5m**).

In summary, MDK exerts repressive roles on DCs at all main levels that define their function: compromising their differentiation but also, blunting their intrinsic ability to uptake antigens, get activated, cross-present antigens to CD8 T cells and promote the cytotoxic activity of these cells.

### MDK reduces the efficacy of DC-based therapies

Given the coordinated effect of MDK on cDCs (differentiation and, function), we anticipated potent inhibitory effects on immunotherapy-based treatments. We tested this hypothesis in the context of anti-DEC-205-OVA vaccination (**Fig. 6a-f**) and in the response to immune checkpoint blockade (**Fig. 6g-h**).

**Figure 6.**
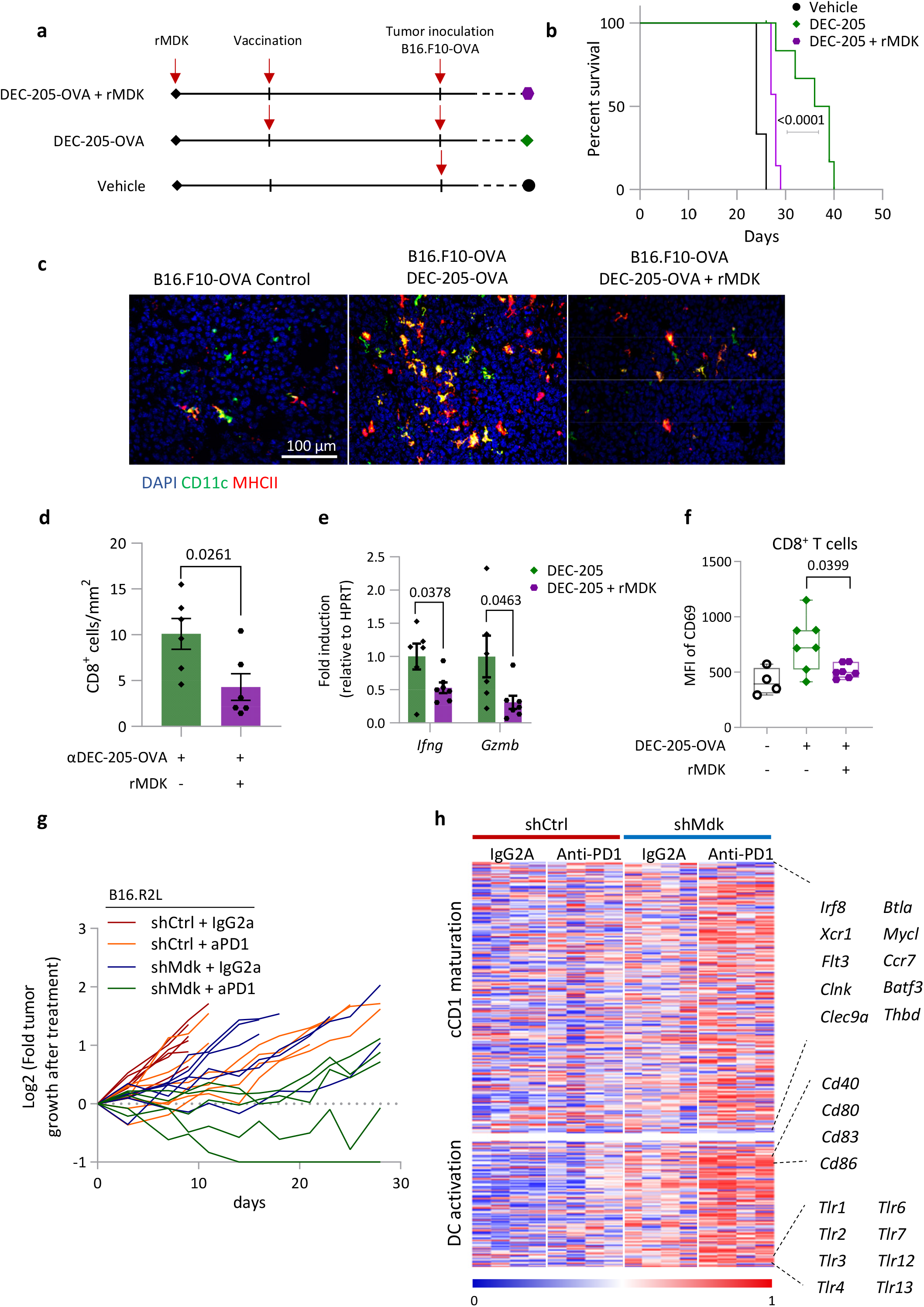
MDK compromises the efficacy of DC-based therapies. **(a)** Schematic representation of the experimental set-up for DEC-205-OVA based treatment. **(b)** Kaplan-Meier curves of overall survival of B16.F10-OVA inoculated in mice not vaccinated (vehicle, black), vaccinated with DEC-205-OVA (green), or vaccinated with DEC-205-OVA + rMDK (purple). Log-rank (Mantel-Cox) test was performed to compare survival among groups (P-value; ***=p<0.001). Sample sizes: no vaccine control n = 3; n = 7 DEC-205-OVA vaccination; n = 7 DEC-205-OVA vaccination + rMDK. **(c)** Immunofluorescence images of intratumoral cDCs (CD11c^+^MHCII^+^: yellow) in B16.F10-OVA tumors from mice immunized with DEC-205-OVA or DEC-205-OVA + rMDK or only PBS. **(d)** Quantification of intratumoral CD8^+^ cells mice vaccinated as indicated. Statistical significance was determined by an unpaired two-tailed t-test. **(e)** Fold induction of gene expression of *Ifng* and *Gzmb* in whole tumors from DEC-205-OVA and DEC-205-OVA + rMDK immunized mice. Statistical significance was determined by unpaired two-tailed t-test. **(f)** Expression of CD69 on CD8^+^ T cells in the blood of mice immunized as indicated (day 20) represented by mean fluorescence intensity (MFI). Statistical significance was determined by unpaired two-tailed t-test. **(g)** Fold tumor growth (Log_2_) curves of B16.R2L-shCtrl or –shMdk implanted in mice and treated with IgG2a isotype control (5 mg kg^-1^) or with anti-PD-1 antibody (clone RMP1-14; 5 mg kg^-1^). Samples sizes were: n = 6 (shCtrl + IgG2a); n = 7 (shCtrl + anti-PD-1); n = 7 (shMDK + IgG2a); and n = 9 mice (shMDK + anti-PD-1). **(h)** Heatmap showing the differential expression of genes from cDC1-related pathways^54^ and DC activation (GO:0001773). Scale bar corresponds to reads per kilobase million (RPKM, log_2_ scale) normalized per gene (0 = min, 1 = max).

Anti-DEC-205-OVA was exploited as an antibody-based strategy to deliver specific antigens (OVA in this case) to cDC1s^58,59^. This antibody was inoculated (with poly(I:C), and the agonist CD40 as a vaccination strategy) 24 h before tumor cell implantation in mice (see schematic in **Fig. 6a**). Parallel treatment arms included animals that were also preconditioned with rMDK, which was also added or omitted to the vaccination mix (**Fig. 6a**). Cells used were B16.F10-OVA (low MDK) to avoid other systemic effects of MDK^17^.

Remarkably, rMDK reduced significantly the anti-tumor response of anti-DEC-205-OVA, resulting in a significantly poorer survival of mice bearing B16.F10-OVA cells (**Fig. 6b**, p<0.0001). Histological analyses confirmed that rMDK nearly abrogated the potent mobilization of CD11c^+^MHCII^+^ DCs by the anti-DEC-205-OVA antibody (**Fig. 6c**, compare right to middle panels). This was translated into a 50% reduction of infiltrated CD8^+^ T cells in the presence of rMDK in the vaccination (**Fig. 6d**). Moreover, RT-PCR analyses showed a reduced expression of genes characteristically associated with T cell cytotoxic activity such as IFN-γ and granzyme B (**Fig. 6e**). By flow cytometry, we found that rMDK reduced the mobilization CD69^+^ CD8^+^ activated T cells by the anti-DEC-205-OVA vaccine (see quantification in the blood of the treated animals in **Fig. 6f**).

Regarding immune checkpoint blockade (ICB), we tested anti-PD1 antibodies, as this treatment has been reported to depend on cDC1s for a proper immune response^48,60^. Consistent with our previous findings^17^, Mdk depletion in melanoma cells improved the antitumor efficacy of anti-PD1 in mouse allographs (**Fig. 6g**). RNAseq showed that the synergistic effect of ICB and Mdk depletion resulted in highly potentiated activation of genes favoring DC activation (GO:0001773) and cDC1 maturation^54^ in the treated allographs (**Fig. 6h**). Therefore, these results support the use of MDK inhibitors to favor cDC mobilization and function in ICB-based treatments.

## Discussion

Here, we report a multi-pronged inactivation of antigen presentation coordinated by MDK that resulted in an immunosuppressive background with implications for patient prognosis and resistance to immune checkpoint blockade. Specifically, we have uncovered a potent suppressive role of MDK on DCs, affecting specially the cDC1 subtypes. Combining RNA sequencing, flow cytometry, histopathological and functional analyses, MDK was found to reduce the amount of DCs and cDC1 in tumors, blood and draining lymph nodes in mouse models. These effects of MDK were supported by an inverse correlation between MDK-and cDC1-associated gene scores in patient datasets (from melanoma and other tumor types). From a mechanistic perspective, our data also show (1) that MDK rewired the transcriptome of DCs, reducing in particular the expression of transcription factors required for the maturation into cDC1. (2) This MDK-mediated control of DC maturation was mediated by STAT3 and traced back to common DC precursors in the bone marrow. (3) MDK-educated DCs displayed significantly impaired functions in antigen uptake, antigen processing, cross-priming and activation of CD8^+^ T cells in vitro and in vivo. (4) Finally, MDK reduced the efficacy of treatments aimed to promote DC-based vaccination and to deactivate immune checkpoint blockers.

Secreted proteins expressed selectively (or preferentially) by tumor cells and impinging on key aspects of cellular proliferation, invasion and metastasis represent attractive targets for drug development. In this context, MDK is raising attention in basic and clinical settings, as this gene is markedly downregulated in adult tissues, but re-expressed in melanoma and a variety of cancer types^26,61,62^. The inverse correlation between MDK- and cDCs-associated scores we identified in patients with melanoma were also demonstrated in adrenocortical carcinoma, lung adenocarcinoma, breast invasive carcinoma, mesothelioma and uterine corpus endometrial carcinoma, expanding the potential therapeutic uses of this factor in oncology. We consider particularly relevant the suppressive roles of MDK in the maturation and mobilization of cDC1 cells. cDC1 are the most potent antigen presenting cells, and their roles on CD8^+^ T activation have a determinant contribution on the antitumoral efficacy of immune checkpoint blockers^48,60^. Moreover, a corollary of our results is that MDK would drive resistance to a variety of treatments in clinical testing amied to target DCs, either by increasing their numbers, delivering specific tumor antigens or promoting a more activated phenotype^38^. Roles of MDK in the polarization of macrophages towards tumor-promoting phenotypes, as well as on the control of pro-metastatic signals via the lymphatic vasculature which we have previously reported, further support pursuing MDK from a pharmacological perspective.

In this study we set to focus on cDC1 based on the coordinated inhibition of key transcription factors traditionally linked to the maturation of this cell type (i.e. *Irf8, Batf3, Id2*, and *Nfil3*) as we demonstrated in bone marrow precursors (**Fig. 4**). The interplay between MDK and cDC1 repression was also reinforced by histopathological studies in mouse allographs and by computational analyses in large clinical datasets we mined with respect to 3 independent cDC1 signatures ^48–51^ However, broader effects of MDK on DC biology are granted given that these are a dynamic and highly complex set of cell types (and states). In particular, it would be interesting to address the role of MDK on mature regulatory DC (mregDCs) which have been recently reported in non-small-cell lung cancer displaying mixed cDC1-cDC2 markers and tolerogenic features^54^. Of note, MDK did not significantly alter cDC2-associated transcription factors such as Irf4 in the BDMC systems we tested in this study. However, MDK-expressing melanoma allographs did decrease the level common DC precursors and pre-DCs at the bone marrow (**Fig. 3h**). Therefore, we expect that MDK may affect different mature DC subtypes in vivo. Curiously, we found cDC2-expressing cells (CD24^-^CD11b^+^CD103^+^) deregulated in mice bearing highly immunogenic B16-R2L melanoma cells, but not in the more immunologically silent B16-F1 counterparts. Dissecting molecular differences between these otherwise isogenic tumor cell variants may pave the way for the identification of autocrine and paracrine signals that may cooperate with MDK in driven immune suppression.

Our studies here have also uncovered gene expression profiles to use as a platform for other cellular environments where MDK may have seemingly opposing roles in immune regulation. To which extent MDK may favor (instead of blocking) DC maturation and function in autoimmune diseases as rheumatoid arthritis, lupus nephritis, and multiple sclerosis^29,31,32^ is worth pursuing. Differential effects on STAT3-associated signaling are granted, considering the negative roles of this transcription factor in the maturation of cDC1s in part via Irf8^63^. Moreover, STAT3 is also well known to regulate other aspects of antigen presentation involving MHC class II and multiple costimulatory molecules, whilst exerting various proinflammatory activities^64^. Context-dependent pro- and anti-tumorigenic roles are also the case for cytokines such as IL-6, TNF-α and NF-κB which are also modulated by MDK.

In summary, here we have identified MDK as a potent immune suppressor in melanoma acting upstream and downstream of DCs: blocking the maturation of precursors at the bone marrow and interfering with all main functions of these cells, from antigen uptake to CD8^+^ T cell-based melanoma cell killing. The signaling cascades reported here may extend to other aggressive tumor types and provide the framework for therapeutic intervention in inflammatory diseases where antigen presentation may be deregulated.

## Abbreviations

BMDCs: bone marrow-derived dendritic cells;
DC: dendritic cells;
cCD1: conventional type I DCs;
GOF: Gain-of-function;
ICB: Immune checkpoint blockade;
LOF: loss-of-function;
LPS: lipopolysaccharide;
MDK: Midkine;
MIF: median fluorescence intensity;
NK: Natural killer cells;
OS: Overall survival;
OVA: ovalbumin;
ssGSEA: single sample gene set enrichment analyses.

---

**Supplementary Figure 1.**
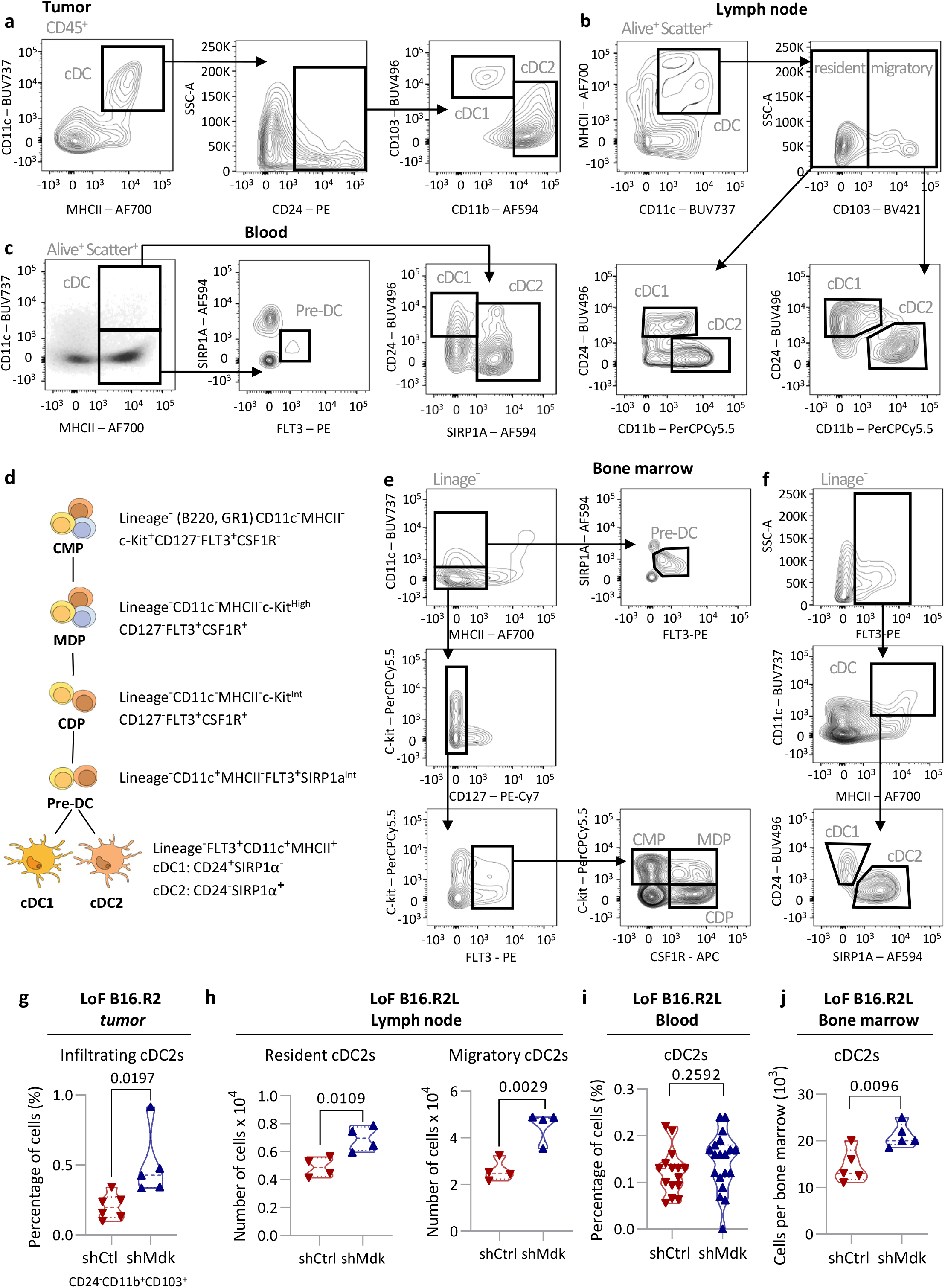
Flow cytometry-based analysis of the impact of MDK on DCs. **(a-c)** Gating strategy to identify intratumoral DCs **(a)**, total cDCs (resident and migratory), DC1s, and DC2s in draining lymph nodes **(b)**, cDCs, DC1s, DC2s, and Pre-DCs in blood **(c)**. **(d)** Schematic drawing of myelopoiesis. **(f)** Gating strategy for bone-marrow identification of pre-DCs, common myeloid progenitors (CMPs), macrophage/dendritic cell progenitors (MDPs) and common dendritic cell progenitors (CDPs), as well as mature cDCs, DC1s, DC2s. **(g)** Infiltrating cDC2 (CD24^-^ CD11b^+^CD103^+^) in tumor implants generated by B16.R2-shCtrl and -shMdk cells (shCtrl: n = 6; shMdk: n =5). **(h)** Resident cDC2 (CD24^+^CD11b^-^) and migratory cDC2 (CD24^-^CD11b^+^) in draining lymph nodes (shCtrl: n = 4; shMdk: n = 4 dLN) of B16.R2L-shCtrl and -shMdk melanomas. **(i)** Percentage of circulating cDC2s in blood of mice bearing B16.R2L-shCtrl (red) vs shMdk (blue) cells (shCtrl: n = 16; shMdk: n =19). Unpaired two-tailed t-test. **(j)** cDC2s in bone marrow (femur and tibia) from mice bearing B16.R2L transduced with shCtrl (red) or shMDK (blue) (shCtrl: n = 5; shMdk: n =5). Statistical significance in panels (g-j was determined by an unpaired two-tailed t-test.

**Supplementary Figure 2.**
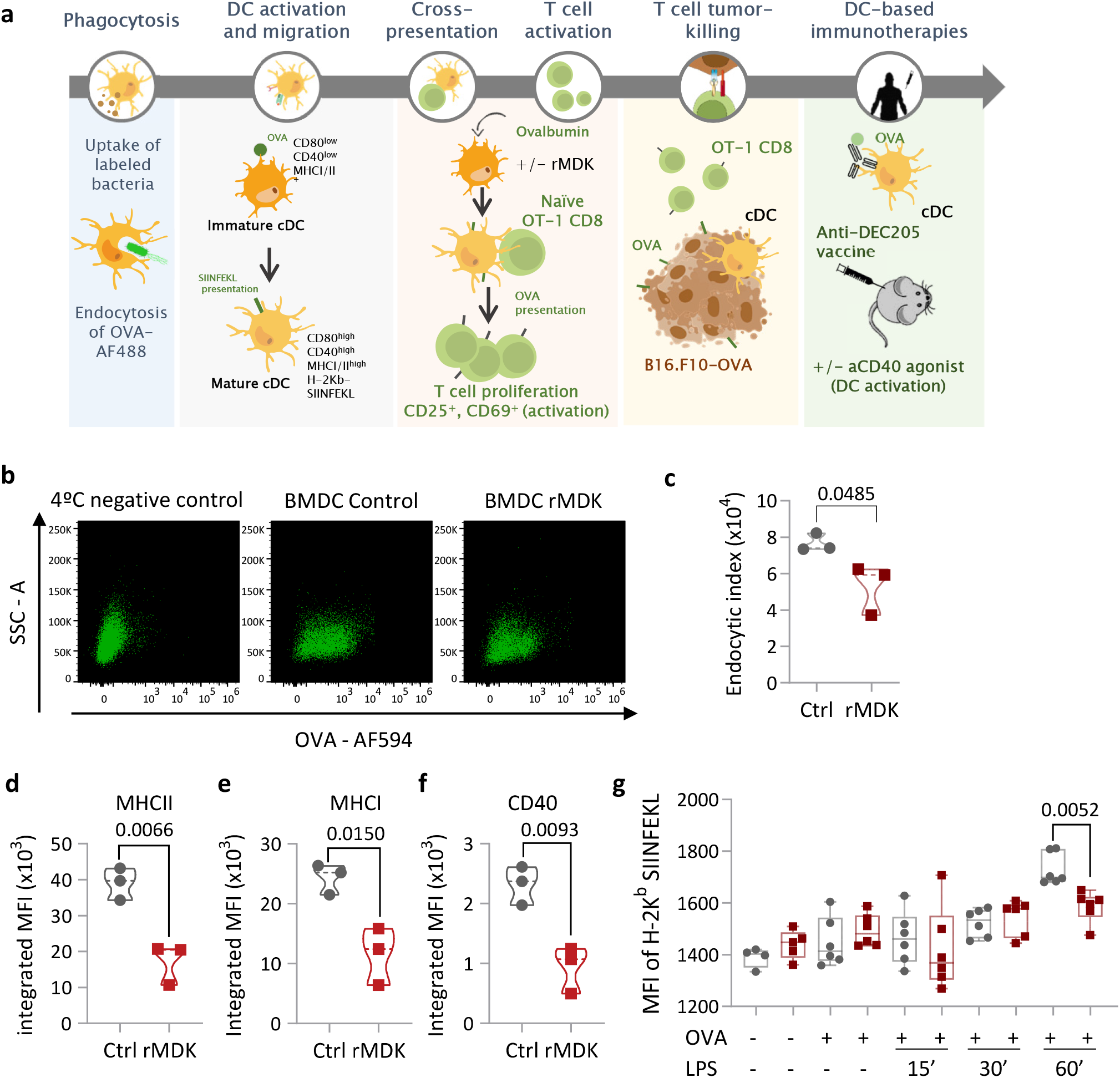
Functional impact of MDK on DC. **(a)** Schematic summary of the methods used to assess DC functions controlled by MDK (see the Methods section for specific detail). **(b)** Dot plot representation of OVA-AF488 endocytosis by BMDCs (CD11c+B220-), in the absence or presence of rMDK (10mg/ml, 48h). Left panels correspond to negative controls of BMDCs kept at 4°C. **(c)** Quantification of endocytosis by cDCs (CD11c^+^B220^-^) without (grey) and with rMDK (red). Endocytic index= (mean fluorescence intensity (MFI) of the phagocytosing cells) x (% fluorescence-positive cells). Statistical significance was determined by unpaired two-tailed t-test. Expression of **(d)** MHCII, **(e)** MHCI, **(f)** CD40 on BMDCs treated with rMDK for 48h defined by flow cytometry. Data are plotted as mean fluorescence intensity (MFI), and the statistical significance was determined by unpaired two-tailed t-test. **(g)** Flow cytometry analysis of H-2Kb-SIINFEKL expression in BMDCs preincubated with rMDK (10 ng/ml, 24h) and then exposed to OVA and LPS (100ng/ml) for the indicated time points. Data are represented as MFI and statistical difference was determined by one-way ANOVA corrected for multiple comparisons (Tukey test).

## Acknowledgments

The authors thank the colleagues at the CNIO Melanoma Group for help and support, including past members Javier Suárez and Susana Frago, which contributed to earlier work in this study. They also thank Isabel Blanco, Soraya Ruiz, Virginia Granda, Sheila Rueda (CNIO), the animal Facility, Histopathological Unit and Confocal Microscopy Unit mouse colonies and histopathological and protein analyses; David Sancho (CNIC; Madrid Spain) for B16-OVA^GFP^ cells and the OT-I mice strain and for scientific guidance and Patrizia Stoitzner (Medical University of Innsbruck) for help with the anti-DEC-205 studies and for advice on analyses of DC activation in vivo. M.S.S. is funded by grants from the Spanish Ministry of Science and Innovation (PID2020-117621RB-I00), Fundación “La Caixa”-Health Research 2019 and 20202 and European Research Council Advanced Grant. D.O. is funded by grants from the Spanish Ministry of Health (AES-PIS PI18/1057) and “Beca Leonardo a Investigadores y Creadores Culturales 2018 de la Fundación BBVA”. X.C. and M.C.A were funded by the Immutrain Marie Skłodowska-Curie ITN Grant.

## Materials and methods

### Melanoma Cells and BMDCs

The murine melanoma cell lines, B16.F1, B16.F10, B16.R2, B16.R2L, YUMM1.1 and YUMM2.1, and the human melanoma cell line WM-164 and SK-Mel-147 were selected for their differential expression of Midkine, distinct metastatic potential, and their genetic background, recapitulating the most frequent genetic alterations in this disease^16,17^. The B16 and the human melanoma cells were cultured in Dulbecco’s modified Eagle’s medium (DMEM, Lonza; BE12-604F/U1) supplemented with 10% fetal bovine serum (FBS, Lonza; DE14-801F). The mouse melanoma cell lines YUMM1.1 and YUMM2.1 were cultured in DMEM-FG12 (1:1) medium with 10% FBS, 1% NEAA (Gibco; 11130-036) and 1% gentamicin (G1264; Sigma–Aldrich). Mycoplasma contamination was regularly tested by quantitative PCR. All of the melanoma cell lines were authenticated using the GenePrint 10 Loci Service.

Bone marrow-derived dendritic cells (BMDCs) were obtained from femurs and tibias of 8-12 weeks-old C57BL/6JOlaHsd mice as reported^39^. In short, bone marrow was extracted by cutting the tips of the bones followed by centrifugation. Next, Erythrocytes Lysis buffer (Qiagen; 79217) was added for 10 min to remove erythrocytes. After lysis, cells were passed through a 40 μm cell strainer. Cells were then plated at 1-1.5×10^6^ cells/ml with 50ng/ml human FLT3-ligand (Miltenyi Biotec; 130-096-480) and cultured for 7 days in RPMI-1640 Glutamax (Gibco; A14517-01) supplemented with 10% heat-inactivated FBS, 1% non-essential amino acids (Gibco; 11130-036), 10 mM HEPES, 1mM Sodium Pyruvate (Gibco; 11360070), 50 nM β-mercaptoethanol, and 50ng/ml gentamicin. On day 7, media supplemented with hFLT3L was renewed for two additional days to fully differentiate BMDCs.

### Gene overexpression and silencing by lentiviral transduction

Overexpression of human MDK was performed using the ORF lentiviral expression vector pReceiver-Lv105-A0792 (MDK) and the corresponding empty vector (Genecopoeia. MDK silencing was performed by lentiviral-driven expression of shRNAs, with pLKO-constructs purchased from Sigma Aldrich as previously reported^16,17,65^. MDK-sh5 (NM_002391.3-621s21c1) was used to deplete MDK in human cells; and for mouse cells, Mdk-sh5 (NM_010784.4-734s1c1) was selected. Non-Target shRNA (CAACAAGATGAAGAGCACCAA) was used as control^16,17,16,17^. Viruses were produced in 293FT cells and infections were performed as previously described, except for B16, where infection included a centrifugation step of 90 min at 1200 rpm. Infected cells were selected by incubation with puromycin (1 μg/ml, Sigma-Aldrich; P-8833) and MDK downregulation or overexpression was determined by ELISA (Peprotech; 900-K190) or qRT-PCR.

B16-OVA cells were generated by lentiviral-mediated transduction of a truncated non-secreted ovalbumin (OVA)-GFP fusion protein (bm1 T OVA) generously supplied by D. Sancho (CNIC, Madrid). B16.F10 cells overexpressing the secreted fraction of the ligand of FLT3 (B16.F10-FLT3LG) were kindly supplied by also by D. Sancho.

### Analysis of DC differentiation and function in vitro

#### Dendritic cell differentiation

BMDCs were differentiated with hrFLT3L as mentioned before. Progenitor cells were treated with 10 ng/ml of hrMDK (PeproTech; 450-16) at days 0, 5 and, 7. MDK effect on BMDCs differentiation was determined by flow cytometry and qPCR. When indicated, BMDCs were cultured with conditioned medium from melanoma cells (ratio 1:1 with fresh complete RPMI).

#### Dendritic cell activation

BMDCs were differentiated with hrFLT3L as mentioned before. Once fully differentiated, 10 ng/ml of hrMDK were added for 48h. Control and MDK-conditioned BMDCs were treated with 100ng/ml of LPS (Sigma; L4391) for 3 h and activation was assessed by flow cytometry, qTR-PCR, or immnoblotting.

#### Endocytosis in dendritic cells

One million of BMDCs cultures in the absence or presence of 10 ng/ml of recombinant MDK (indicate provider and origin) were incubated with 1 μg/ml of Ovalbumin-AF594 conjugate (Thermo Fisher Scientific; O34781) for 30 min at 37°C. A non-endocytosis control was prepared at 4°C. Endocytosis was assessed by cytometry in a BD FACSCanto™ II Cell Analyzer (BD Biosciences). The Endocytosis Index was calculated as the mean fluorescence intensity (MFI) per cell with respect of the percentage of cells that had incorporated the OVA-AF594 conjugate.

#### Phagocytic activity of DCs

BMDCs incubated in the absence or presence or rMDK were stained with DiD’ solid (Thermo Fisher Scientific; D7757) to be traced by fluorescence microscopy in the far-red emission channel. To this end, the corresponding BMDCs were plated at 4×10^5^ cells/ml per well in a μ-Slide 8 Well (IBIDI; 80826) together with 10 μg of pHRodo Green *E. coli* BioParticles (Thermo Fisher Scientific; P35366). BMDCs pre-treated with 1 μM Cytochalasin D (Sigma-Aldrich; C8273), an inhibitor of actin polymerization that blocks phagocytosis, were used as a negative control. The assay was performed on a Leica DMI6000B wide-field fluorescence microscope with an incubation chamber and CO2 control. Images for the red channel (DiD’ solid tracer) and green channel (pHRodo) were captured every 7 minutes with a 20X objective (HCXPLAPO 0.7 NA). Image analysis was done by using a customized routine programmed on ImageJ software. This macro segmented the cells by using the red channel and then quantified the green signal for every frame. Fluorescence intensity was averaged for each cell and normalized with respect to the first time frame.

#### Antigen presentation and antigen-specific tumor cell killing

Control- or MDK-conditioned BMDCs were pulsed with 250 ng/ml of chicken ovalbumin (OVA, grade IV, Sigma) overnight. The next day, spleens from OT-I mice were disaggregated and erythrocytes were removed. Splenocytes were counted and CD8^+^ T cells were sorted with the LS MACS columns (Miltenyi Biotech; 130-042-401) using the mouse CD8 (Ly-2) MicroBeads (Miltenyi Biotech; 130-117-044). After sorting, CD8^+^ T cells were stained with 5 μM of CellTrace Violet Cell Proliferation Kit (Thermo Fisher Scientific, C34557), following the manufacturer’s instructions. Then, BMDCs were co-cultured for 3 days with OT-I T cells at a ratio of 1:1 (500,000 cells) in a 96 well-rounded plate. Clonal T cell proliferation was analyzed by flow cytometry on a BD LSRFortessa Flow Cytometer (BD Biosciences). These OT-1 cells activated by BMDCs conditioned by MDK were then cocultured with B16.F10-OVA (expressing GFP). T cell killing was assessed by DAPI and Annexin V-APC labeling in GFP positive melanoma cells. also using a BD LSRFortessa Flow Cytometer. When indicated, BMDCs were treated with the STAT3 inhibitor Silibinin (1 μM, Sigma-Aldrich; S0417)), or DMSO for 3 h prior addition of rMDK.

### *In vivo* assays

#### Murine strains

C57BL/6JOlaHsd mice were purchased from Harlan or bred at the CNIO mouse facility. The *Batf3* knockout mice (*Batf3*^tm1Kmm^)^44^ and OT-I mice (C57BL/6JOlaHsd-*Ptprc^a^Rag1^ko/ko^Tg(TcraTcrb)1100Mjb/J)^66^* developed by Dr. Michael J. Bevan (University of Washington Medical Center, USA) were kindly provided by D. Sancho (CNIC, Madrid).

#### Generation of melanoma allographs

7-9-week-old mice were used as recipients for subcutaneous (s.c.) injection of 5×10^5^ murine melanomacells (of the indicated backgrounds expressing or lacking MDK). Tumor growth was followed every two days by measuring the two orthogonal external diameters using a caliper. Tumor volume was calculated using the formula (a x b^2^ x 0.52)^1^. Mice were sacrificed at the human endpoint when tumor volume reached 1 cm^3^. All experiments with mice were performed in accordance with protocols approved by the Institutional Ethics Committee of the CNIO and the Instituto de Salud Carlos III.

#### Vaccination and DC activation

Vaccination of 7-9-week-old mice (C57BL/6JOlaHsd) was performed by s.c. administration of 5 μg of chicken ovalbumin (OVA, grade IV, Sigma Aldrich; A5503-1G) mixed with the adjuvants anti-CD40 (12.5 μg, Clone 3/23, BD Biosciences; 553787) and poly(I:C) HMW (12.5 μg; InvivoGen; TLRL-PIC) in PBS (total volume of 50 μl). When indicated, hrMDK (1 μg) was added to the vaccination cocktail. 3 days after vaccination, mice were sacrificed and activation of DCs in draining inguinal lymph nodes was assessed by flow cytometry as indicated below.

#### Antigen presentation

Naïve CD8^+^ T cells isolated from OT-I mice (considering that mice need to be from the same gender as the vaccinated mice) were obtained and stained with 5 μM of CellTrace Violet Cell Proliferation Kit. These labeled OT-I cells were injected retro-orbitally (10^6^ cells in 100 μl) on mice vaccinated as indicated above (OVA/anti-CD40/pI:C). 3 days afterwards, inguinal lymph nodes were harvested to analyze OT-I T cell proliferation by flow cytometry (as the reduction in CellTrace Violet signal).

#### Prophylactic DEC-205 vaccination

For this protocol, mice were pre-conditioned by s.c injection of 2 μg of hrMDK (in 50 μl of PBS). 24 h afterwards, a second s.c. was performed with 0.5 μg anti-DEC-205-OVA or IgG-OVA (kindly provided by Diana Dudziak), these reagents were mixed with 25 μg anti-CD40 and 25 μg poly(I:C) HMW, with or without 2 μg of hrMDK. 7 days after vaccination 5×10^5^ B16.F10-OVA cells were injected s.c. injected. Tumor growth was followed every two days.

#### Treatment with immune checkpoint blockers

Mice inoculated s.c. with 5×10^5^ B16.R2L-shMdk or their isogenic shControl were treated with the anti-PD1 agonist antibody (clone RMP-14; BioXCell) or its IgG2a control isotype (clone 2A3; BioXCell). Treatment was performed intraperitoneally every 3 days at a dose concentration of 10 mg kg^-1^ body weight.

### RNA extraction and RT-PCR

Total RNA was extracted and purified from cell pellets using QIAshredder and RNeasy^®^ Mini-Kit (QIAGEN; 79654 and 74104) following the manufacturer’s instructions, and RNA concentration was determined by NanoDrop Spectrophotometer ND-1000 (NanoDrop Biotechnologies). In the case of whole tissue RNA extraction, tissues were stored in RNA Later solution (Invitrogen; AM7020) for 24h at 4°C and later stored at −80°C. Tissue (30 mg) was then homogenized with 1.4 mm ceramic beads (Qiagen; 13113-325) and lysed with RLT buffer from RNeasy Mini-Kit in a Precellys 24 (Bertin Technologies). QIAshredder and RNeasy Mini-Kit and where then used for RNA isolation.

Next, cDNA was produced by reverse-transcribing 1 ug of total RNA using the SuperScript III Reverse Transcriptase (Thermo Fisher Scientific; 18080093), according to the manufacturer’s protocol. A total of 40 ng of cDNA was used as a template for the qPCR reaction. Finally, real-time quantitative polymerase chain reaction (RT-qPCR −60°C annealing temperature) was performed with *Power* SYBR Green PCR Master Mix (Applied Biosystems; 4367659). Assays were run in triplicates on the QuantStudio 6 Flex Real-Time PCR System (Applied Biosystems). Primers were designed with ROCHE Tool and are listed in **Table 1**. HPRT and 18s were used to normalize mRNA expression.

**Table 1.**
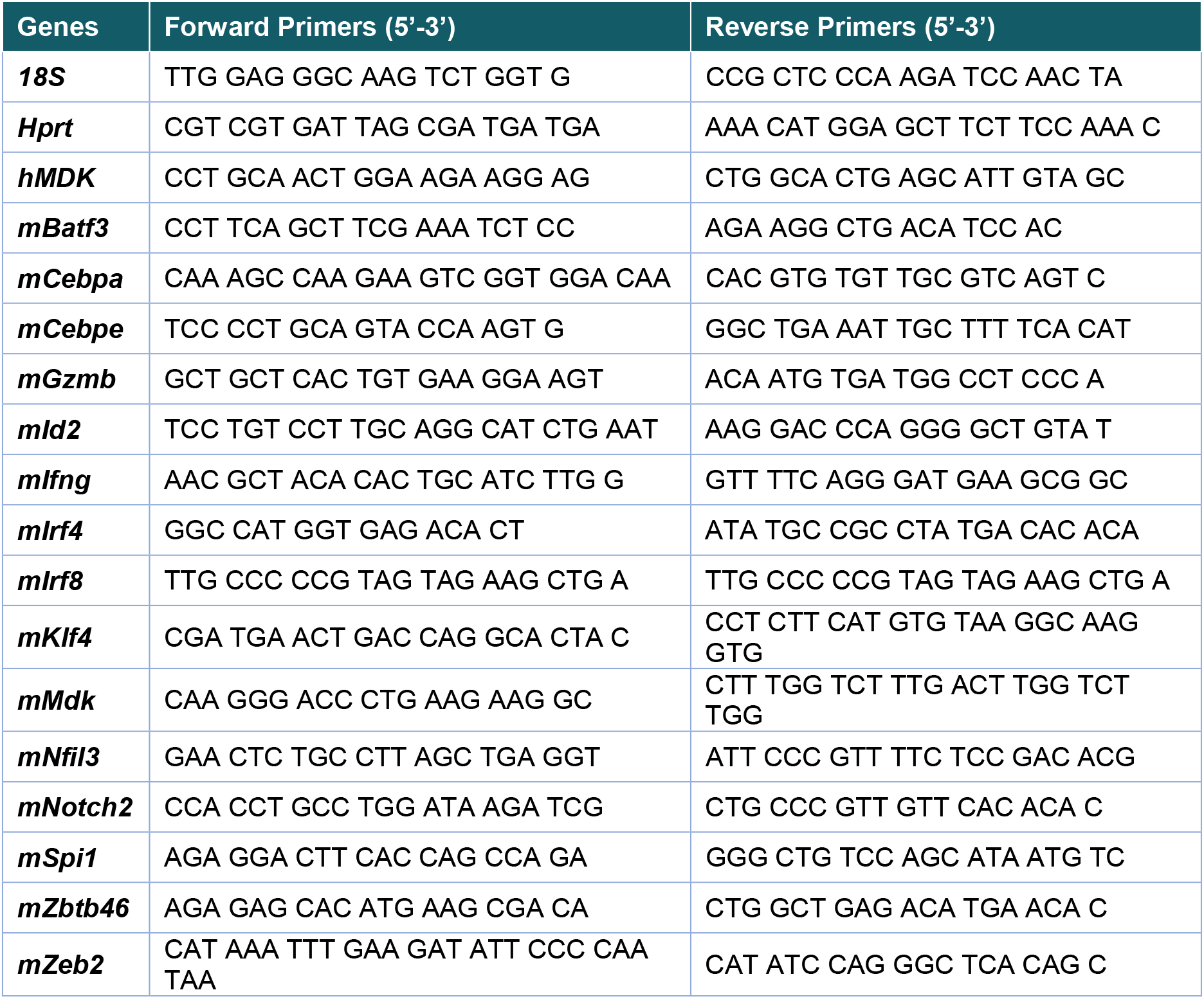
List of primers for RT-QPCR.

### Flow cytometry

#### Sample (tissue) preparation

Tumors were dissected from mice and the total weight of the tumor was determined. Tumors were then minced using scalpels and digested with 500 U/ml Collagenase IV (Gibco; 17104-019), and 200 mg/ml DNAse I (Roche; 10104159001) per 0.3 grams of tumor weight (30 min, 37°C). Samples were then passed through a 40 μm cell strainer (Falcon; 352340) to remove large pieces of undigested tissue.

Lymph nodes were dissected and all fat was carefully removed. Disaggregation was performed on RPMI-1640 with an enzyme mix comprised of 0.8 mg/ml Dispase (Roche; 4942078001), 0.2 mg/ml Collagenase P (Roche; 1121385700) and 0.1 mg/ml DNase I (Roche; 10104159001). First, lymph nodes were minced and incubated for 20 min at 37°C in a water bath, gently inverting every 5 min. Media with the cell suspension was placed in a tube containing ice-cold FACS buffer (DPBS (Gibco; 14190-094), 2% Bovine serum albumin (BSA, Sigma-Aldrich; A7906) and 5 mM EDTA (ED141C)). If lymph nodes were not completely dissociated, the enzymatic cocktail was refreshed, and samples were again incubated at 37°C, while gently pipetting, until complete disaggregation.

Bone marrow was extracted as in BMDCs differentiation protocol. Peripheral blood, blood was drawn from the mouse cheek or from the heart of sacrificed mice using with a 30-gauge syringe pre-wet with 0.5M EDTA. 100 μl of blood were used for downstream staining, with prior erythrocyte lysis.

#### Marker detection and quantification by flow cytometry

For cell surface staining, 1-5 x10^6^ cells, depending on the cell population of interest and organ assessed, were incubated with anti-Fc receptor blocking antibody (clone 2.4G2; BD Pharmingen) in FACS buffer for 30 min on ice. Cells were then stained with the indicated antibodies (see Table 2) in FACS buffer for 30 min on ice. Viability staining was performed with DAPI, or in the case that cells were going to be fixed afterwards, with Zombie Aqua Fixable Viability Kit (Biolegend; 423102) before the blocking. Fixation was performed with 4% paraformaldehyde or Foxp3/Transcription Factor Staining Buffer Set (Thermo Fisher Scientific; 00-5523-00) following the manufacturer’s instructions. Flow cytometry was performed on a BD LSRFortessa Flow Cytometer or BD FACSCanto II Cell Analyzer (BD Biosciences). The following antibodies were used:

**Table 2.** List of antibodies used in flow cytometry.

| Target | Fluorophores | Clone | Reference | Company |
| --- | --- | --- | --- | --- |
| **Annexin V** | APC |  | 550474 | Pharmingen |
| **CD3** | AF700 | 17A2 | 56-0032-80 | Invitrogen |
| **CD4** | PE-Cy7 | RM4-5 | 552775 | BD Pharmingen |
| **CD8a** | FITC | 53-6.7 | 553030 | BD Pharmingen |
| **CD8a** | APC-Cy7 | 53-6.7 | 100713 | BioLegend |
| **CD11b** | PerCP-Cy5.5 | M1/70 | 550993 | BD Pharmingen |
| **CD11c** | BUV737 | HL3 | 564986 | BD Pharmingen |
| **CD16/32** |  | 2.4G2 | 553142 | BD Pharmingen |
| **CD24** | PE | M1/69 | 561079 | BD Pharmingen |
| **CD24** | BUV496 | M1/69 | 6129553 | BD Pharmingen |
| **CD25** | PE-Cy5 | PC61 | 102010 | BioLegend |
| **CD40** | PE-Cy7 | 3/23 | 124621 | BioLegend |
| **CD45** | PE Texas Red | I3/2.3 | ab51482 | Abcam |
| **CD45.1** | PE-Cy7 | A20 | 110729 | BioLegend |
| **CD45R (B220)** | AF700 | RA3-6B2 | 557957 | BD Pharmingen |
| **CD45R (B220)** | PE-Cy7 | RA3-6B2 | 103221 | BioLegend |
| **CD69** | APC | H1.2F3 | 104513 | BioLegend |
| **CD80** | BUV563 | 16-10A1 | 741272 | BD Pharmingen |
| **CD103** | BV421 | 2E7 | 121421 | BioLegend |
| **CD115 (CSF1-R)** | APC | AFS98 | 135509 | BioLegend |
| **CD117 (c-Kit)** | PerCP-Cy5.5 | 2B8 | 105823 | BioLegend |
| **CD127 (IL7R)** | PE-Cy7 | A7R34 | 135013 | BioLegend |
| **CD135 (FLT3)** | PE | A2F10 | 135305 | BioLegend |
| **CD197 (CCR7)** | PerCP-Cy5.5 | 4B12 | 120115 | BioLegend |
| **CD279 (PD-1)** | BV711 | 29F.1A12 | 135231 | BioLegend |
| **Gr1** | PE-Cy7 | RB6-8C5 | 552985 | BD Pharmingen |
| **H-2K<sup>b</sup> bound to SINFEKL** | PE | 25-D1.16 | 141603 | BioLegend |
| **MHCI (H-2K<sup>b</sup>)** | APC | AF6-88,5 | 116517 | BioLegend |
| **MHCII (I-A/I-E)** | AF700 | M5/114.15.2 | 107622 | BioLegend |
| **NK-1.1** | BUV421 | PK136 | 562921 | BD Pharmingen |
| **OVA** | AF594 |  | O34783 | ThermoFisher |
| **Sirp1a (CD172a)** | FITC | P84 | 144005 | BioLegend |
| **Sirp1a (CD172a)** | AF594 | P84 | 144020 | BioLegend |
| **Phospho-Stat3 (Tyr705)** | AF647 | O4-46 | 4324 | BD Horizon |
| **XCR1** | PE | REA707 | 130-111-372 | Miltenyi Biotec |

Analyses of flow cytometry data were done using Flowjo (Treestar). Cell populations were discriminated after gating on single cells (discriminated by FSC-A and FSC-W), excluding non-viable cells and removing debris (discriminated by FSC-A and SSC-A) using the following combinations of cell markers:

**Table 3.**
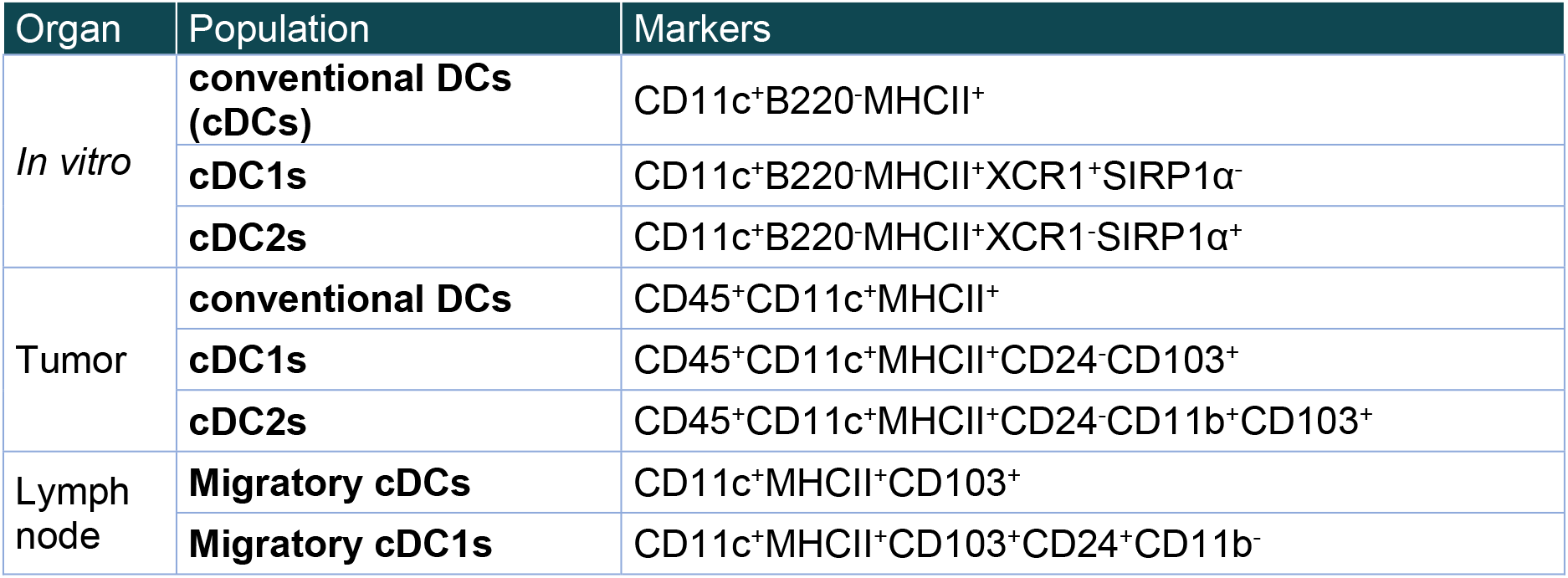
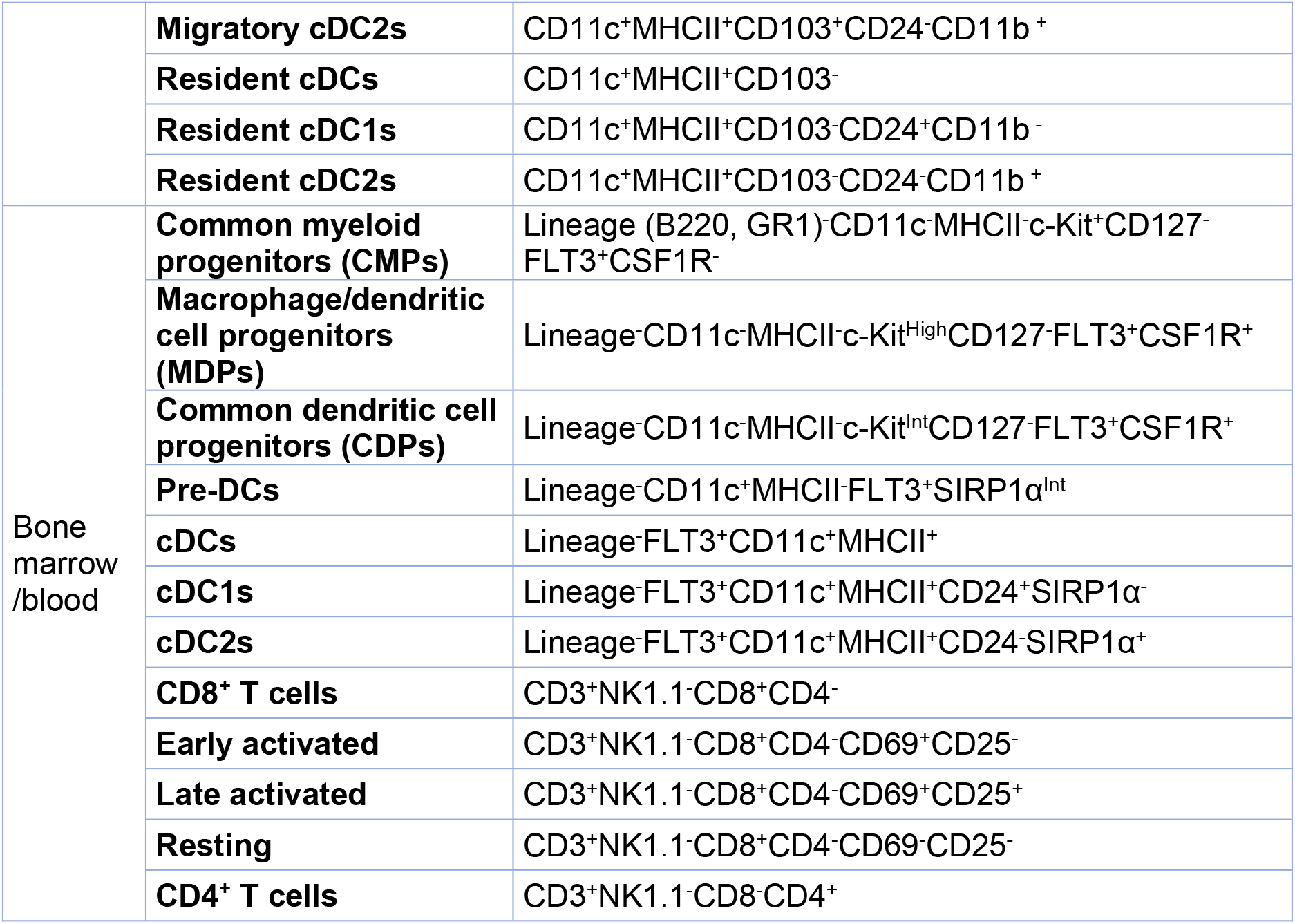
Gating strategies used in flow cytometry analysis.

### Histological analyses

#### Staining by immunofluorescence

Tissues were embedded and frozen on OCT compound (Tissue-Tek, 4583) and cut with a Microm HM 550 OVPD Cryostat (Thermo Fisher Scientific; 956464). Samples were then fixed with the Antigenfix buffer (Diapath, P0016) for 15 min at RT. Following, samples were permeabilized with PBS Tween-20 0.02% and 0.4% Triton-100 for 1 h. Next, tissues were blocked with PBS 3% BSA Tween-20 Normal Goat serum for 30 min. Antibodies used where: CD11c-AF488 (1:100; N418 Biolegend; 117313) and MHCII (1:300; M5/114.15.2; Biolegend; 107601)). Primary antibodies were incubated overnight at 4°C in a humidified chamber. After washing with PBS 3 times, samples were stained with the corresponding secondary antibody at 1:1000, depending on the species of the primary antibody. Nuclei were counterstained with DAPI. Negative controls were obtained by omitting the primary antibody. Images were acquired in a TCS SP5 AOBS (Acousto Optical Beam Splitter) confocal microscope (Leica). Images were subsequently analyzed with ImageJ software.

#### Staining by immunohistochemistry

Tissues were fixed in formalin and embedded in paraffin. The immunohistochemistry reactions were performed in an automated immunostaining platform (AS Link, Dako, Agilent) by the Histopathology Unit at CNIO. In brief, antigen retrieval was first performed with the appropriate buffer, (Low pH buffer) and endogenous peroxidase was blocked (peroxide hydrogen at 3%). Then, slides were incubated with rat monoclonal anti-CD8 (OTO94A, CNIO Monoclonal Antibodies Core Unit). After the primary antibody, slides were incubated with the corresponding secondary antibodies and visualization systems (Novolink Polymer Linker, Leica) conjugated with horseradish peroxidase. The immunohistochemical reaction was developed using 3, 30-diaminobenzidine tetrahydrochloride (DAB), and nuclei were counterstained with Carazzi’s hematoxylin. Next, slides were mounted with a permanent mounting medium for microscopic evaluation. Positive control sections known to be primary antibody positive were included for each staining run. Whole slides were captured with a slide scanner (AxioScan Z1, Zeiss), and images captured with the Zen Blue Software (V3.1 Zeiss).

### RNA sequencing

RNA sequencing was performed by Genomics Unit at CNIO. The RNA integrity was evaluated by Agilent 2100 Bioanalyzer using RNA 6000 Pico kit following the manufacturer’s recommendations. In brief, sequencing libraries were prepared with the QuantSeq 3’ mRNA-Seq Library Prep Kit FWD for Illumina (Lexogen; catalog number 015) following the manufacturer’s instructions. Complementary DNA (cDNA) libraries were purified, applied to an Illumina flow cell for cluster generation, and sequenced on an Illumina NextSeq 550 (with version 2.5 reagent kits) following the manufacturer’s protocols. Single-end sequenced reads followed adapter and polyA tail removal as indicated by Lexogen. The resulting reads were analyzed with the nextpresso pipeline as follows. Sequencing quality was checked with FastQC version 0.11.0. Reads were aligned to the mouse genome (GRCm38/mm10) with TopHat 2.0.10 using Bowtie 1.0.0 and SAMtools 0.1.19, allowing three mismatches and 20 multihits. The GENCODE vM20 gene annotation for GRCm38 was used. Read counts were obtained with HTSeq. Differential expression and normalization were performed with DESeq2, filtering out those genes where the normalized count value was <2 in more than 50% of the samples.

#### Bioinformatic analyses

mRNA profiles generated by RNAseq as indicated before in sc implants of the indicated cell lines were processed to identify significantly enriched pathways (p≤0.05) using Cytoscape v3.8.2 and the ClueGO v2.5.7 plug-in, focusing on “Gene Annotation (GO) immune system process”. Gene Set Enrichment Analysis (GSEA) was performed using annotations from libraries of molecular Signature Databases (MSigDB) and Gene Annotation databases. After Kolmogorov-Smirnoff correction for multiple testing, only those pathways bearing an FDR <0.25 were considered significant^97^. Heatmaps were created by Morpheus (log_2_ normalization).

#### ssGSEA and scores generation

For assessing MDK relevance in clinical data, the top upregulated genes (top 100-upregulated and bottom 44-downregulated genes) from RNA-seq of B16.R2L-shMdk vs –shCtrl allografts were used for single-sample Gene Set Enrichment Analysis (ssGSEA) (https://www.genepattern.org/). This allowed to generate an MDK-in vivo associated score (MDK-iS) in biopsies of metastatic melanomas from The Cancer Genome Atlas (TCGA). TCGA melanoma samples whose mRNA expression profiles scored above or below the 33th percentile in this MDK-iS score were selected for subsequent analysis of expression profiles and overall survival. This cut-off was selected as it allowed for a sizable number of biopsies to represent patients with the most separate MDK-associated mRNA expression profiles.

RNA-seq was also performed on BMDCs treated or not with rMDK. Data was then used for ssGSEA to define an MDK-educated DCs score in biopsies of primary and metastatic melanomas from TCGA. Melanoma samples whose mRNA expression profiles scored above or below the 20th percentile were chosen for downstream analyses.

